# Phase variation as a major mechanism of adaptation in *Mycobacterium tuberculosis* complex

**DOI:** 10.1101/2022.06.10.495637

**Authors:** Roger Vargas, Michael J. Luna, Luca Freschi, Kenan C. Murphy, Thomas R. Ioerger, Christopher M. Sassetti, Maha R. Farhat

## Abstract

Phase variation induced by insertions and deletions (INDELs) in genomic homopolymeric tracts (HT) can silence and regulate genes in pathogenic bacteria but this process is not characterized in MTBC adaptation. We leverage 31,428 diverse clinical isolates to identify genomic regions including phase-variants under positive selection. Of 87,651 INDEL events that emerge repeatedly across the phylogeny, 12.4% are phase-variants within HTs (0.02% of the genome by length). We estimated the *in-vitro* frameshift rate in a neutral HT at 100x the neutral substitution rate at 1.1 × 10^−5^ frameshifts/HT/year. Using neutral evolution simulations, we identified 4,098 substitutions and 45 phase-variants to be putatively adaptive to MTBC (P<0.002). We experimentally confirm that a putatively adaptive phase-variant alters the expression of *espA,* a critical mediator of ESX-1 dependent virulence. Our evidence supports a new hypothesis that phase variation in the ESX-1 system of MTBC can act as a toggle between antigenicity and survival in the host.

## INTRODUCTION

Tuberculosis (TB), caused by pathogens of the *Mycobacterium tuberculosis* complex (MTBC), is a major public health threat causing an estimated 10 million new cases of disease per year (World Health Organization, 2020). Human TB is primarily caused by seven major phylogenetic lineages (L1-L7) also known as *M. tuberculosis sensu stricto*, and two more distant human-adapted MTBC lineages L5, L6 are also known as *M. africanum* (Gagneux, 2018). More recently, studies have revealed two new lineages: L8 in Uganda and Rwanda (Ngabonziza et al., 2020) and L9 in East Africa (Coscolla et al., 2021).

MTBC genomes show no evidence for recombination or horizontal gene transfer. Genomic diversity, including more ancient divergence from the MTBC ancestor and between lineage members, is instead driven predominantly by DNA damage and replication error resulting in chromosomal point mutations. A different mechanism with 100-1000 fold faster kinetics is the development of insertion and deletions in short sequence repeats (SSRs) of 1-7bp nucleotides through mispairing (Van Der Woude and Bäumler, 2004). This slipped-strand mispairing (SSM) occurs with misalignment between repeats on the mother and daughter strands during DNA synthesis resulting in an increase or decrease in the number of repeat units in the newly synthesized strand (Van Der Woude and Bäumler, 2004). These changes can result in frameshifts or alteration in a transcriptional regulatory region leading to phase-variable expression of a protein. Repeats of a single nucleotide, or homopolymeric tracts (HTs), is the simplest form of SSR. SSM within these regions was recently observed in the MTBC resulting in antibiotic tolerance or resistance (Bellerose et al., 2019; Safi et al., 2019; Vargas and Farhat, 2020).

Of the variants generated by mutation or SSM, the vast majority do not reach appreciable population allele frequencies. The allele frequency spectrum in MTBC supports a high proportion of low-frequency variants, especially singletons consistent with background and/or purifying selection on average across the genome (Gagneux, 2018; Pepperell et al., 2010). In specific regions, variants may arise more than once in parallel (*i.e.*, among bacterial strains that do no share an immediate common ancestor). This is rare under neutral theory or purifying selection but can be observed due to population demographic shifts or due to positive selection (Brynildsrud et al., 2018; Farhat et al., 2013). Parallel evolution has been commonly observed in antibiotic resistance genes and specifically variants that allow the organism to withstand antibiotic killing (Farhat et al., 2013; Gagneux, 2018; Manson et al., 2017). More recently, parallel evolution has been observed in connection with enhanced virulence and transmission (Brynildsrud et al., 2018; Chiner-Oms et al., 2019; Holt et al., 2018; Vargas et al., 2021).

Here, we leverage a sample of 31,428 geographically diverse clinical isolates that have undergone whole-genome sequencing (WGS) and are representative of the genetic diversity found within the MTBC. These isolates represent more than 30,000 natural evolution experiments of MTBC infecting humans and transmitting to the next host. Using data on these isolates, we infer the number of times each genetic variant has evolved in a parallel fashion within and outside of HTs in the MTBC genome. With simulations, we determine which variants are likely under positive selection. Using precise genome engineering, we functionally validate HT variants measured to be under positive selection that occur in a regulatory region of the MTBC virulence factor *espA*, a gene essential for type VII ESX-1 mediated secretion. The results support that MTBC continues to evolve towards a phenotype of more effective patient-to-patient transmission.

## RESULTS

### Genetic diversity in 31,428 MTBC clinical isolates

We curated and processed 33,873 publicly available genomes. For quality control, we excluded 1,663 isolates with inadequate sequencing data at ≥ 10% of variable sites curated across the full dataset (**Figure S1A**-**S2, Materials and Methods**). We excluded an additional 290 isolates because they could not be typed into an MTBC major lineage based on SNV barcode (most commonly because of missing calls at lineage defining sites, **Materials and Methods, Fig. S2**) (Freschi et al., 2021); excluded 35 isolates because they belonged to L7 that was otherwise not well represented, and excluded 457 isolates because they were typed into L4 but not an L4 sub- lineage, the latter needed for computational efficiency of the phylogeny estimation (**Fig. S2**). In the remaining 31,428 isolates, we detected 836,901 single nucleotide variants (SNV) occurring at 782,565 genomic sites across the 4.4-Mb MTBC genome (17.7%) (**Figure S1A**-**S2**, **Materials and Methods**). Of the 782,565 SNV sites, 422,891 (54.04%) were singletons, *i.e.* only a single isolate harbored a minor allele at that site. Additionally, we detected 47,425 INDELs with 27,937 (58.9%) being singletons (**Figure S1A**, **Materials and Methods**).

For computational efficiency, the 31,428 isolates phylogeny was constructed separately for L1, L2, L3, L4 (split into three subgroups L4A,B,C), L5 and L6 (**Figure S3, Figure S1A-S2, Materials and Methods**) (Edwards et al., 2020). We built a multiple sequence alignment of SNV sites and used maximum-likelihood phylogenetic estimation. The phylogenies represented well the global *M. tuberculosis sensu stricto* diversity: spanning 2,815 isolates from L1, 8,090 L2, 3,398 L3, 5,839 L4A, 6,958 L4B, 4,134 L4C; *M. africanum* was represented by 98 L5 and 96 L6 isolates. The SNV barcode misclassified only 14/31,428 isolates compared with the full phylogenetic reconstruction (**Materials and Methods**). Given the size of the phylogeny that challenged visualization, we computed t-Distributed Stochastic Neighbor Embeddings (t-SNE) of the matrix of pairwise SNP distances (**Figure S4A**, **Materials and Methods**). We visualized isolates in this t-SNE embedding space labeling isolates by lineage and confirmed good separation between sub- lineages especially at short scales (**Figure S3**, **S4B-I**). Within-lineage diversity was congruent with expected diversity, including highest diversity within L1, L4 and L6 and lowest diversity within L2 (**Figure S4J**) (Coscolla and Gagneux, 2014).

### Parallel evolution

Using maximum likelihood ancestral reconstruction, we computed the number of parallel/repeated arisals of minor allele SNV mutations (homoplasy score or Hs) across the eight phylogenies (**Figure S3**, **S5A-B**, **Materials and Methods**). As ancestral reconstruction methods cannot infer INDEL events simultaneously with SNVs, we developed an alternative method (TopDis) to assess separately for INDEL parallel evolution. TopDis relies on observing monophyletic groups harboring the derived allele that are separated in the tree by isolates harboring the reference state (**Figure S5C**, **Materials and Methods**). We confirmed the accuracy of the TopDis approach by computing TopDis Hs for SNVs and showing they are equal to Hs computed using ancestral reconstruction for most variants (**Figure S1B-C**, **Figure S6**).

### Putatively adaptive SNVs

The distribution of homoplasy scores for SNVs was strongly right skewed; 102 SNVs were acquired ≥ 100 times (**Materials and Methods**, **Figure 1A**, **Table S1**, **Table S2**) (Manson et al., 2017). Population bottlenecks can increase the rate of parallel evolution observable in a phylogeny, but estimates of effective population size for Mtb over similar time and geographic scales, which have been modeled with constant and exponential growth priors, did not identify evidence for population contraction (O’Neill et al., 2019). Mtb molecular clock rate estimates have also been robust to assumptions of constant *vs.* exponential population growth under a coalescent model (Menardo et al., 2019). Here, to simulate the expected rate of parallel mutation acquisition under neutral evolution, we ran simulations using a range of estimated molecular clock rates for *M. tuberculosis* assuming a constant population size (**Materials and Methods**) (Menardo et al., 2019). We estimated SNVs to arise with Hs ≥ 5 with probability <0.002 under these assumptions. In our data Hs ≥ 5 was observed for 4,980 (0.49%) of SNV sites (**Figure 1**). Of the subset of 1,525/4,980 with a minor allele frequency >0.1% (**Figure 1C-D**), 470 (30.8%) were coding synonymous, 738 (48.4%) were coding non-synonymous, 308 (20.2%) were intergenic, and 9 (0.59%) were in non-coding RNA regions. Sites in genomic regions associated with antibiotic resistance represented 13 of the top 30 sites by Hs (>222) (**Figure 1D**, **Table S1**, **Table S2**).

**Figure 1.**
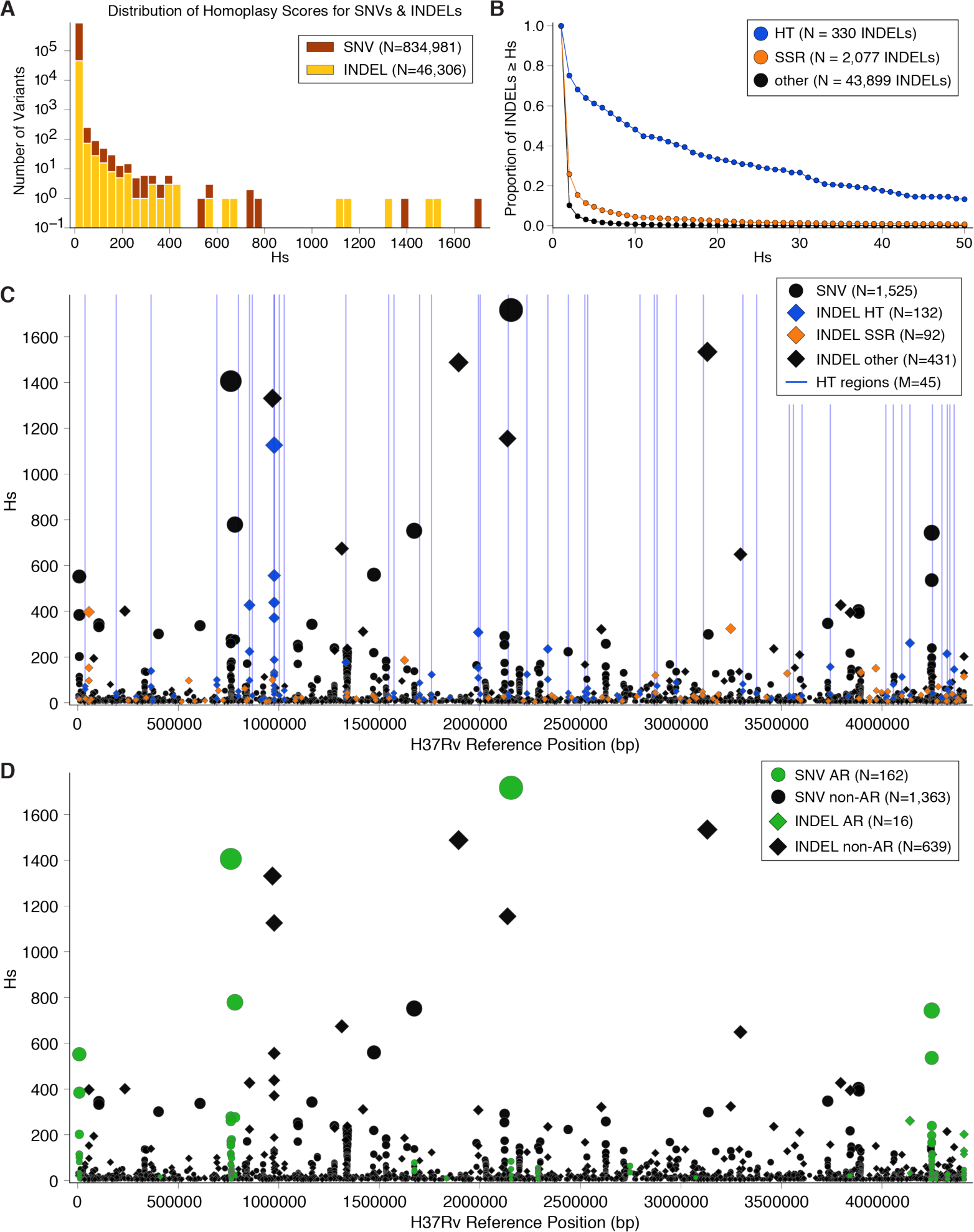
Parallel evolution of SNVs and INDELs. (**A**) The distribution of homoplasy scores for 834,981 SNVs and 46,306 INDELs. 0.49% of SNVs have a homoplasy score ≥ 5 (*P* < 0.002) and 3.01% of INDELs have a homoplasy score ≥ 5. (**B**) Proportion of INDELs with Hs ≥ *x* for varying values of *x*, split into sets according to whether INDEL occurs within HT, SSR or other region of the genome. (**C**-**D**) Homoplasy score (Hs) for 1,525 SNVs and 655 INDELs with homoplasy score ≥ 5 and minor (SNVs)/alternate (INDELs) allele frequency > 0.1% among 31,428 isolates, plotted against position on the genome. Bubble size corresponds to Hs. (**C**) INDELs broken down by whether they occur within an HT, SSR or other region of the genome. HTs with a cumulative Hs score > 45 (across INDELs occurring within HT) are indicted by blue bars. (**D**) Variants colored in green occur within loci that have been associated with Antibiotic Resistance.

**Table 1.** HTs with significant association with antibiotic resistance. Associations between frameshift variants in HTs and resistance to antibiotics. Variants in 17 HT regions were significantly associated with resistance to at least one antibiotic at the Bonferroni corrected threshold (**Methods**). S: number of isolates susceptible, R: number of isolates resistant, FS: number of isolates that harbor a frameshift, WT: number of isolates with wild type state. *For HTs associated with resistance to more than one antibiotic, details for the most significant association are reported while other antibiotics are listed in the last column. AMK: Amikacin, CAP: Capreomycin, CYS: Cycloserine, EMB: Ethambutol, ETA: Ethionamide, INH: Isoniazid, KAN: Kanamycin, MXF: Moxifloxacin, OFX: Ofloxacin, PZA: Pyrazinamide, RFB: Rifabutin, RIF: Rifampicin, STR: Streptomycin

| Gene Symbol | HT H37Rv<br>coords | Hs | drug | S<br>(FS/WT) | R<br>(FS/WT) | OR 95% CI<br>(Fisher Exact Test) | -log10(bonf<br>p-val) | *other antibiotics |
| --- | --- | --- | --- | --- | --- | --- | --- | --- |
| *Rv2264c* | 2536625-2536632 | 138 | STR | 4341<br>(968/3373) | 2101<br>(985/116) | 29.59 (24.1-36.33) | 90.3 | AMK, CAP, EMB, INH, KAN,<br>MXF, OFX, PZA, RFB, RIF |
| *lysX-infC* | 1852176-1852183 | 29 | MXF | 3243<br>(22/3221) | 338<br>(49/289) | 24.82 (14.8-41.64) | 67.6 | AMK, CAP, CYS, EMB, ETA,<br>INH, KAN, OFX, PZA, RFB,<br>RIF, STR |
| *glpK* | 4139183-4139190 | 282 | RIF | 10.89k<br>(50/10840) | 3868<br>(172/3696) | 10.09 (7.35-13.85) | 66.6 | EMB, ETA, INH, PZA, RFB,<br>RIF, STR |
| *Rv3413c* | 3832356-3832363 | 39 | KAN | 3077<br>(5/3072) | 577<br>(35/542) | 39.68 (15.48-101.72) | 34.0 | AMK, CAP, EMB, INH, RIF,<br>STR |
| *Rv2177c-aroG* | 2440187-2440194 | 69 | PZA | 9018<br>(174/8844) | 1804<br>(121/1683) | 3.65 (2.88-4.64) | 27.5 | CAP, EMB, ETA, INH, RIF |
| *PE_PGRS25* | 1572680-1572687 | 68 | PZA | 9018<br>(345/8673) | 1804<br>(178/1626) | 2.75 (2.28-3.32) | 25.2 | EMB, INH, RIF |
| *bioF2* | 36470-36477 | 140 | EMB | 9307<br>(1395/7912) | 2394<br>(558/1836) | 1.72 (1.54-1.93) | 19.8 | INH, PZA, RFB, RIF |
| *vapC2-Rv0302* | 364498-364505 | 216 | PZA | 9018<br>(153/8865) | 1804<br>(97/1707) | 3.29 (2.54-4.27) | 18.8 | EMB, ETA, INH, RIF |
| *Rv1373* | 1546465-1546472 | 58 | RIF | 10890<br>(204/10686) | 28<br>(21/3847) | 0.29 (0.18-0.45) | 6.2 | CFX, EMB, INH, MXF, RIF |
| *Rv3192* | 3559990-3559997 | 55 | EMB | 9307<br>(25/9282) | 2394<br>(31/2363) | 4.87 (2.87-8.27) | 8.1 | RIF |
| *Rv0759c-Rv0760c* | 854252-854261 | 776 | EMB | 9307<br>(8483/824) | 2394<br>(2272/122) | 1.81 (1.49-2.2) | 6.8 | AMK, CAP, INH, KAN, PZA,<br>RIF |
| *lipR* | 3450182-3450189 | 40 | EMB | 9307<br>(18/9289) | 2394<br>(21/2373) | 4.57 (2.43-8.58) | 4.7 | INH |
| *Rv1894c* | 2141408-2141415 | 72 | RFB | 431<br>(5/426) | 607<br>(43/564) | 6.5 (2.55-16.54) | 3.3 | PZA |
| *Rv0694-Rv0695* | 794672-794679 | 8 | MXF | 3243<br>(0/3243) | 338<br>(2/336) | N/A | 3.0 |  |
| *espK* | 4358979-4358986 | 192 | PZA | 9018<br>(177/8841) | 1804<br>(65/1739) | 1.87 (1.4-2.49) | 2.8 | INH, MXF |
| *PE_PGRS31* | 2001789-2001796 | 57 | INH | 9.844k<br>(79/9765) | 4693<br>(72/4621) | 1.93 (1.4-2.66) | 2.4 |  |
| *mshD-phoT* | 912694-912701 | 27 | CAP | 3611<br>(0/3611) | 663<br>(3/660) | N/A | 2.4 |  |

### Homopolymer tracts demonstrate a high concentration of INDELs

Because INDELs can be generated by SSM or other mutational processes depending on the genetic sequence context, we divided the 46,306 observed INDELs into the following groups: (1) INDELs in HT regions (n=330 in 145 unique HTs), (2) INDELs in more complex SSR of a pattern of 2 to 6 base-pairs (bp, n=2,077), and (3) INDELs in other regions of the genome (n=43,899) (**Figure 1B**, **Figure S7, Materials and Methods**). In HTs, the INDEL acquisition rate across the phylogeny normalized by aggregate region length was 9,339.7/kbp, compared to 61.8/kbp in other SSR regions and 16.9/kbp elsewhere on the genome (*P* < 1*x*10^−100^ across three tests for difference between Poisson rates). For comparison, the SNV acquisition rate across the genome was 242.6/kbp aggregated from 834,981 SNVs detected genome-wide. Further, 75.2% of the INDELs in HT regions were homoplastic at a score Hs > 1 compared to 25.9% and 10.3% of INDELs called in all SSR regions and non-HT-SSR, respectively (**Figure 1B-C**, **Figure S7**).

### Putatively adaptive INDELs

Of the 46,306 total INDELs observed, the majority or 32,883 (71%) caused frameshifts within open reading frames with a median allele frequency across the sample of 0.003%. The distribution of INDEL acquisitions across the phylogeny was strongly right skewed with 59 mutations acquired independently ≥ 100 times (**Figure 1A**, **Table S3**, **Table S4**). Compared with SNVs, a higher relative proportion of INDELs demonstrated Hs ≥ 5 (1,393/46,306, 3.01%, P-value < 1x10^−5^, Fisher Exact test) (**Figures 1A**, **1C**). Of the 655/1,393 INDELs with allele frequency >0.1%, 132/655 (20.1%) were in HT regions and 94/352 (26.7%) of the subset that resulted in frameshifts occurred in HT regions (**Figure 1C**, **Table S4**). A lower proportion of INDELs was found in known antibiotic resistance-associated genes compared to SNVs (16/655 vs 162/1525, P-value = 7x10^−11^, Fisher Exact test) (**Figure 1D**, **Table S2**, **S4**). Among the 30 INDELs with the highest Hs (>187), only three occurred in genes associated with antibiotic resistance: *gid* 103delC (Hs = 202) did not occur within an SSR or HT region and is known to confer streptomycin resistance (**Table S3**) (Coll et al., 2018; Manson et al., 2017), *glpK* nt565-572insC (Hs = 261) located within an HT region and previously implicated in multi-drug tolerance (**Figure 1D**, **Table S3**) (Bellerose et al., 2019; Safi et al., 2019), and *ponA1* nt1878insCCGCCGCCT (Hs = 397) located within an SSR region in a gene that contributes to peptidoglycan biosynthesis and alters sensitivity to the antibiotic rifampicin (Farhat et al., 2013).

Given differences in mutational processes and rates at SSR versus other sites, we studied potentially adaptive INDELs separately by whether or not they occur in SSRs. We used a Hs cutoff of ≥ 5, similar to SNVs above. Of the 43,899 non-SSR INDELs, 993 (2.3%) demonstrated an Hs ≥ 5 (**Figure 1B**). The INDEL with the highest Hs was a three amino-acid insertion in the putative antigenic protein Rv2823c that was acquired independently 1,534 times affecting 5,093 isolates across members of the six lineages we evaluated (**Table S3**). The INDELs with Hs ≥ 5 were more likely to affect intergenic regions than INDELs with Hs < 5 (257/993, 26% vs. 6,941/42,906, 16%, P-value = 1.5x10^−14^, Fisher Exact Test).

While intragenic SSM often introduce frameshifts and disrupt ORFs, phase variation at intergenic sites can also have important effects on gene expression (Van Der Woude and Bäumler, 2004). We compared the general features of intragenic phase variation with INDELs that putatively alter gene expression based on their occurrence within 50bp upstream of MTBC transcriptional start sites (Shell et al., 2015) and within regulatory non-coding RNAs (Gerrick et al., 2018) (2,077 SSR INDELs and 330 HT INDELs). Overall, we identified frameshift INDELs in HT and other SSR (294/330, 89.1% of HT INDELs and 1,190/2,077, 57.3% of other SSR INDELs) in open reading frames. Of non-HT SSR INDELs, 6.2% (128/2,077) putatively affect gene expression, and 47.2% (981/2,077) introduce translational frameshifts (**Figure 1B-C, Figure S8**). A greater proportion of INDELs in HT regions were found in likely regulatory regions 7.6% (25/330) and open reading frames 69.7% (230/330) compared to other SSR INDELs (**Figure 1B-C, Figure S7**). The majority of frameshifting INDELs incur a premature stop codon within the first 3/4th of a gene (570/981, 58.1% for SSR INDELs and 117/230, 50.9% for HT INDELs) (**Figure S7**).

Given the measured high rate of frameshift INDELs in HT regions, the expected rapid kinetics of SSM, and the high rate of INDEL homoplasy across the genome, we experimentally measured the neutral rate of +1 frameshifting in a 7G HT derived from the *glpK* gene in *M. smegmatis*. The measured rate was 3.14 × 10^−8^ frameshifts/generation (**Materials and Methods**). Assuming that MTBC doubles once per day on average, this corresponds to a rate of 1.14 × 10^−5^ frameshifts/ HT/year [lower bound = 7.96 × 10^−6^, upper bound = 1.49 × 10^−5^] (**Materials and Methods**). To identify potentially adaptive INDELs that should demonstrate more extreme homoplasy than observed under neutral evolution, we ran simulations of HT evolution respecting the 8 observed Mtb phylogenies (**Figure S8**, **Materials and Methods**). We estimated the probability of HT accumulating >45 INDELs across the phylogeny at <0.002 under the neutral rate (**Figure S8**). Forty-five HTs had a homoplasy score > 45 (**Figure 1C**, **Table S5, Table S6**). These putatively adaptive HTs occurred in one aforementioned gene associated with antibiotic resistance, *glpK*, and the remaining were in other genes spanning a range of functions. Two of the three HT regions with the highest Hs occurred in the 3’ end of *ppe13* (*Hs* = 2,317 and *Hs* = 771), and located 15bp from the stop codon on the 1,332bp ORF (**Table S3, S5**). Of the 3,088 mutation arisals within these adjacent HTs, 49.5% (1,529/3,088) resulted in a premature stop codon while 50.5% (1,559/3,088) resulted in an aberration of the stop codon in the annotated H37Rv gene sequence. Further, 10/45 (22%) of the putatively adaptive HTs occurred in intergenic regions and of these 3/10 occurred within 50bp upstream of a TSS (*Rv3848-espR*, *vapC2-Rv0302*, *espA-ephA*).

### Recency estimation of putative adaptive variants

We hypothesized that if positive selection is driving parallel evolution of an allele then the ratio of homoplasic instances of that allele divided by the number of isolates carrying the same allele can capture the recency of positive selection. We separated genes into four non-redundant categories: *antigen* genes, *antibiotic resistance* genes, *PE/PPE*, and other genes (**Materials and Methods**). We compared other categories to antibiotic resistance genes, as the selection pressure on variants in the latter only commenced with the introduction of antibiotics for Mtb treatment 70-80 years ago (Ektefaie et al., 2021). We computed a recency ratio (RcR) for the 1208 homoplastic SNVs in coding regions. The RcR displayed a strongly right-skewed distribution as most SNVs have very few independent arisals relative to the number of isolates that harbor the minor allele indicating older selection (**Figure 2A**-**B**, **Table S1, Table S2**). As expected, RcR values were highest (indicating more recent evolution) for SNVs in antibiotic resistance regions (*P* < 1 × 10^−16^, Mann-Whitney U-test between antibiotic resistance and every other gene category) (**Figure 2C**).

**Figure 2.**
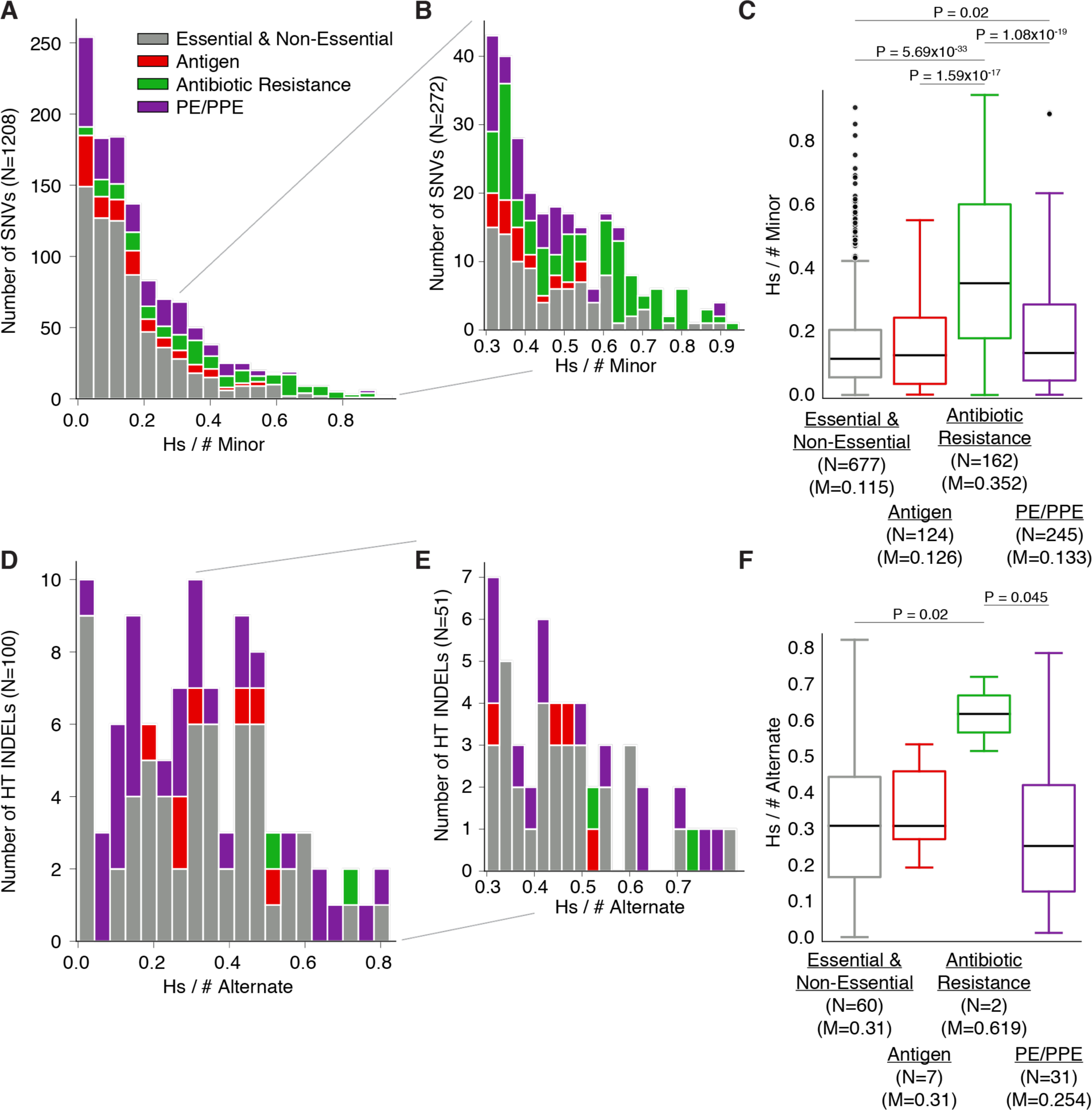
Recency Ratio for SNVs and HT INDELs. (**A-B**) The distribution of the ratio of (homoplasy score) to (# of isolates harboring the minor allele) for 1,208/1,525 SNVs (Figure 1C) that occur in coding regions. (**C**) Breaking these SNV recency ratios down by gene category reveals higher ratios overall for antibiotic resistance genes when compared to other gene categories. (**D-E**) The distribution of the ratio of (homoplasy score) to (# of isolates harboring the alternate allele) for 100/655 INDELs (Figure 1C) that occur in HT and coding regions. (**F**) Breaking these INDEL ratios down by gene category reveals higher ratios overall for antibiotic resistance genes when compared to other gene categories, however the only two INDELs in this gene category were found in the HT of *glpK*. N = number of alleles, M = median RcR

The RcR for the 388 coding non-HT INDELs (grouping non-HT SSR and non-SSR INDELs together) closely resembled that for SNVs (**Figure S9A-B**). Similar to SNVs, RcR values for non-HT INDELs were higher in antibiotic resistance regions (*P* < 0.002, Mann-Whitney U-test between antibiotic resistance and every other gene category) and median RcR values within gene categories mirrored those for observed for SNVs (**Figure S9C**). This suggests that the mutational or other processes giving rise to non-HT INDELs and selection on them is similar to SNVs.

The RcR distribution for the 100 coding HT INDELs demonstrated a shift toward higher values than SNVs or non-HT INDELs in every gene category (**Figure 2D-F**, **Table S3**, **Table S4**). As INDELs in SSR are uniquely prone to revert to the ancestral sequence, this observation may be related to recent selection for the derived allele, recent selection for reversion to the ancestral allele, or both. Regardless, this observation implies recent selection for INDELs in HT tracts.

### Frameshifts in a HT upstream *espA* alter transcription

To assess the functional consequence of variation we observed in HTs (**Figure 1B-C**, **Table S5**), we carried out a genome-wide association with the antibiotic resistance phenotype to 15 antibiotics to uncover any previously unknown associations between frameshift mutations in HTs and resistance to a panel of antibiotics (n= 101-14,537, **Materials and Methods**). Of the 145 HTs studied, 17 were significantly associated with resistance to at least one antibiotic, including the previously known association between convergent frameshifts in the HT of *glpK* and multi-drug resistance (**Figure 3A-B**, **Table 1**). In addition to *glpK*, frameshifts in the HT of Rv2264c (Hs=138) and *lysX-infC* (Hs=29) were the top three positively associated HTs with multi-drug resistance. The majority of HTs (128/145, 88%) do not, however, appear to potentiate antibiotic resistance. We hypothesized that these regions may be mediating a different form of pathogenic adaptation.

**Figure 3.**
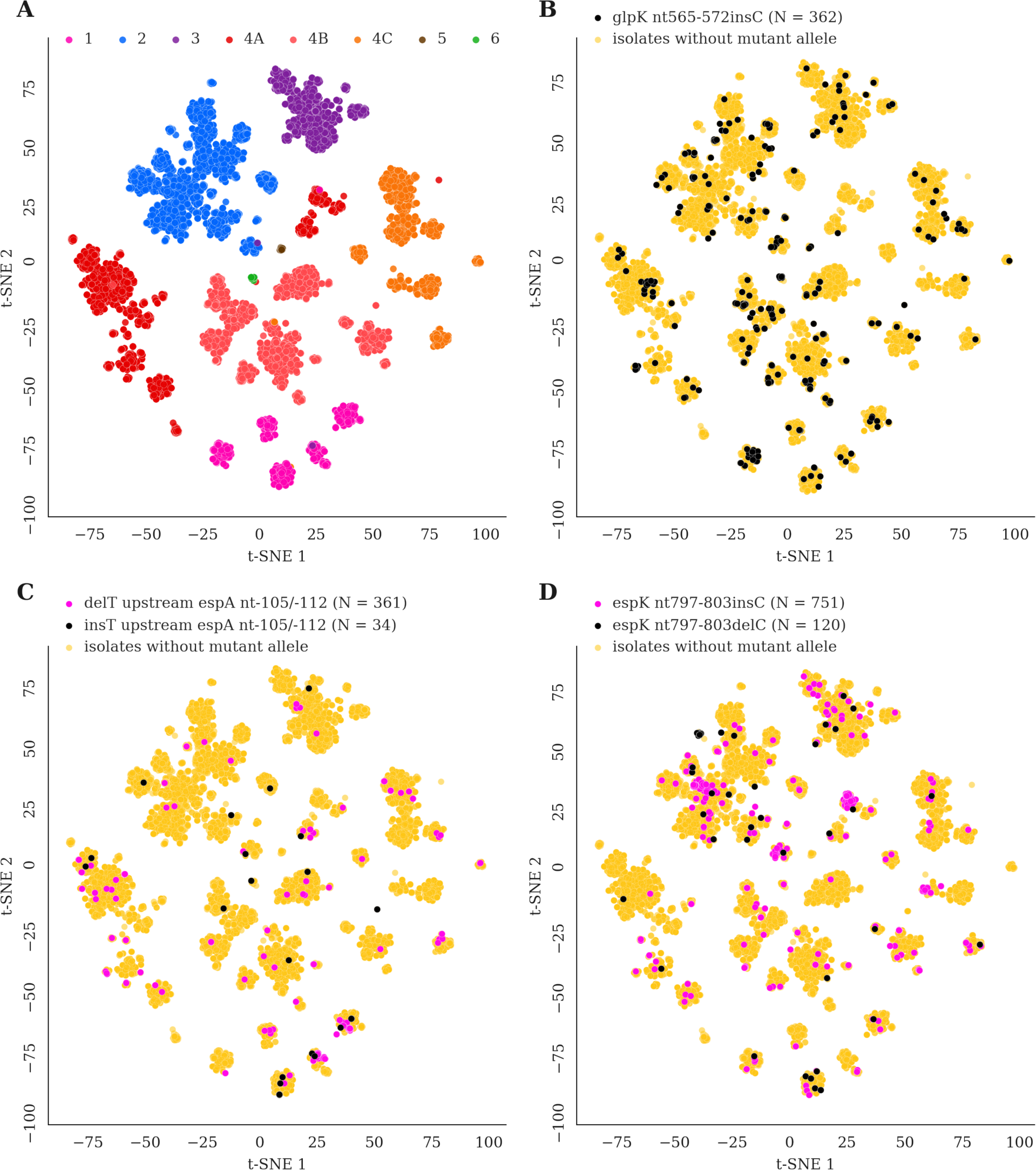
Genetic map confirms homoplastic variants. (**A**) The t-SNE plot serves as a genetic similarity map, isolates are colored according to which group they belong to (L1, L2, L3, L4A, L4B, L4C, L5, L6). (**B-D**) Isolates are labeled if they harbor a given mutant allele (N = # of isolates that harbor the mutant allele). These mutations within HTs (*glpK* nt565-572insC, delT upstream *espA* nt-105/-112, insT upstream *espA* nt-105/-112, *espK* nt797-803insC and *espK* nt797-803delC) are detected in isolates belonging to different clusters, confirming that these mutations must have arisen independently in different genetic backgrounds.

As mentioned above our top HT and non-HT INDEL hits occurred in *PPE13*, and in a putatively antigenic protein respectively suggesting that they mediate adaptation at the immune or host-pathogen interface. The PPE13 HT frameshifts are predicted to shorten the protein product by ∼5 AA, and hence were difficult to evaluate experimentally. We noted that other HTs with high Hs appeared in or near ESX-1 related genes (**Tables S5**). These regions include: (1) The HT between *espA* and *ephA* (ESX1 components that control the rate of secretion) is optimally suited to act as a UP element as a poly-A stretch found ∼48bp upstream of one of two putative transcriptional start sites of the *espACD* operon, (Estrem et al., 1999) (**Figure 4A**), ESX1 components that control the rate of secretion (**Figure 3C**), (2) An intragenic HT disrupts the open reading frame of the ESX1- associated *espK* gene (**Figure 3D**), and (3) An HT in the 5’ UTR of the ESX1 regulator, *espR* (**Table S5**). To assess the phenotypic consequence of these mutations, we engineered the most abundant +1 HT variant upstream *espACD* operon into the H37Rv genome and assessed the effect of this variant on gene regulation during exponential growth in 7H9 broth (**Figure 4A**). Comparing the transcriptome of this mutant to its isogenic parent, we found only a small number (22) of significantly differentially expressed genes, most prominently a decrease in the expression of *espA*, *espC*, and *espD* (by approximately 40%, log2-fold-changes=-0.7) (**Figure 4B-C**), along with the downstream genes Rv3613c and Rv3612c (**Table S7**). These data verify the functional effect of this intragenic HT INDEL and suggest positive selection for decreased ESX1 activity.

**Figure 4.**
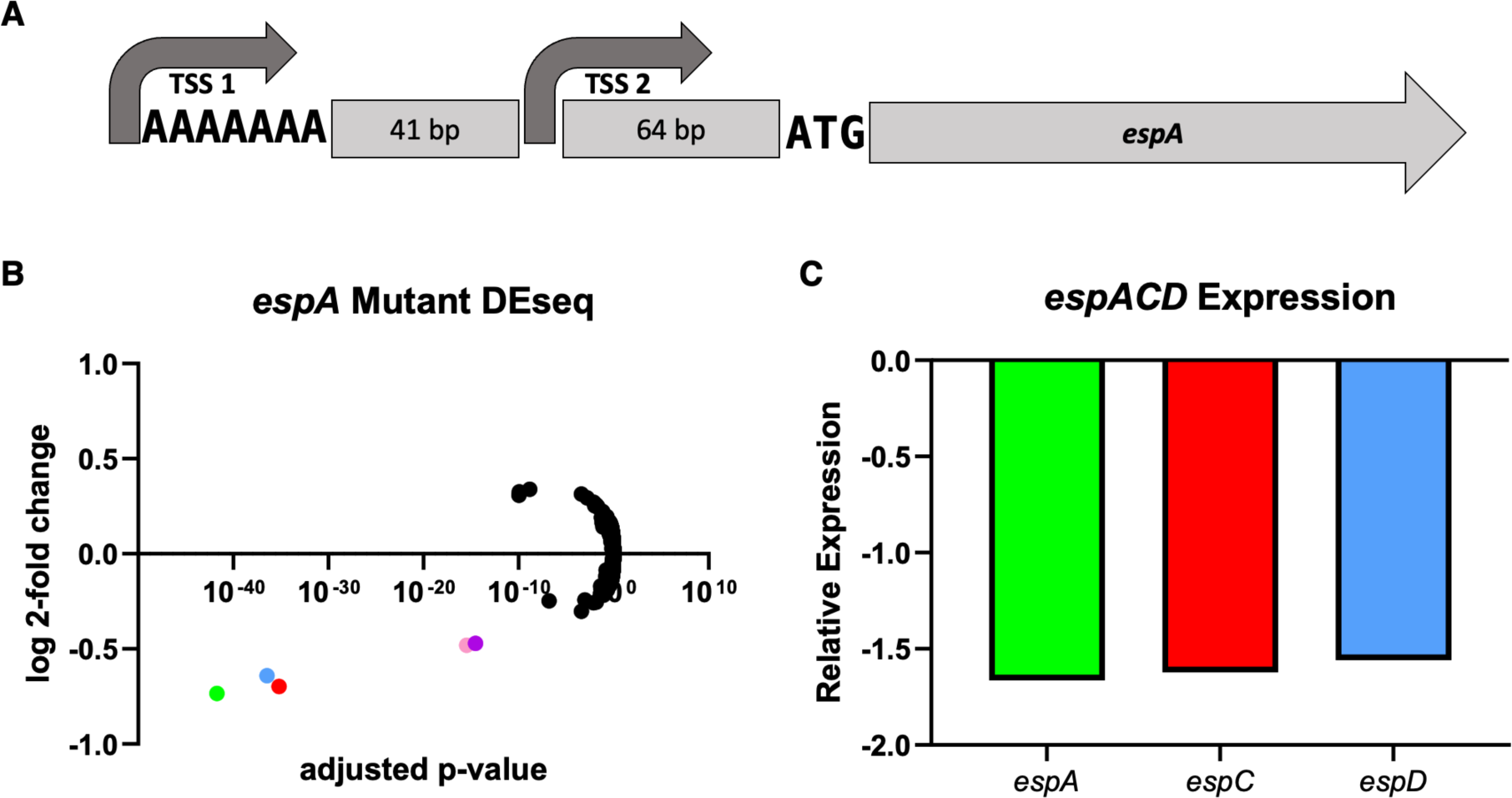
A single basepair deletion within the *espA* homopolymer results in decreased *espA* expression. (**A**) Schematic showing location of 7 basepair homopolymer Upstream of *Rv3616c*. A highly variable, 7 basepair adenine repeat 105 basepairs upstream of the translational start site for *Rv3616c* (*espA*), which forms an operon with downstream genes *espCD*. Upstream of *Rv3616c*, two transcriptional start sites have been identified. The longer of which sits along the homopolymeric stretch, the other is found another 41 basepairs downstream of the homopolymer. A single basepair deletion in the poly-A tract results in a ∼2-fold decrease in *espACD* expression. (**B**) A volcano plot highlighting the results of an RNAseq experiment comparing a recombineered *espA* homopolymer mutant to WT H37Rv. Results are pooled from 2 independent experiments consisting of at least 3 biological replicates each. *espA* (green), *espC* (red), and *espD* (blue) are highlighted. Also highlighted *Rv3612c* (purple) *and Rv3613c* (pink), two genes immediately downstream of *espACD*. (**C**) Relative expression levels of the *espACD* operon in the mutant *espA* strain compared to WT H37Rv.

### Gene-wide mutational density reveals variable ESX and PE/PPE genes

Given the apparent convergence of HT variants on ESX-1 function, we aggregated independent variant arisals at the gene-level to better understand the adaptive landscape of genomic variants in MTBC. Specifically, we aggregated Hs for all variants found within each gene (regardless of frequency) and normalized the resulting score by gene length to obtain the mutational density (**Materials and Methods**). We separated this analysis by SNVs (**Figure 5A**, **Table S8**) and INDELs (**Figure 5B**, **Table S9**) because Hs were computed differently for each (**Materials and Methods**), and because of the different mechanisms at play in generating each type of diversity. We simulated the number of arisals that occur on each gene using a modified molecular clock rate normalized by gene length to obtain a neutral mutation rate for each gene (**Materials and Methods**). We found that a gene has an estimated neutral mutational density ≥ 0.45 with probability <0.002 under these assumptions.

**Figure 5.**
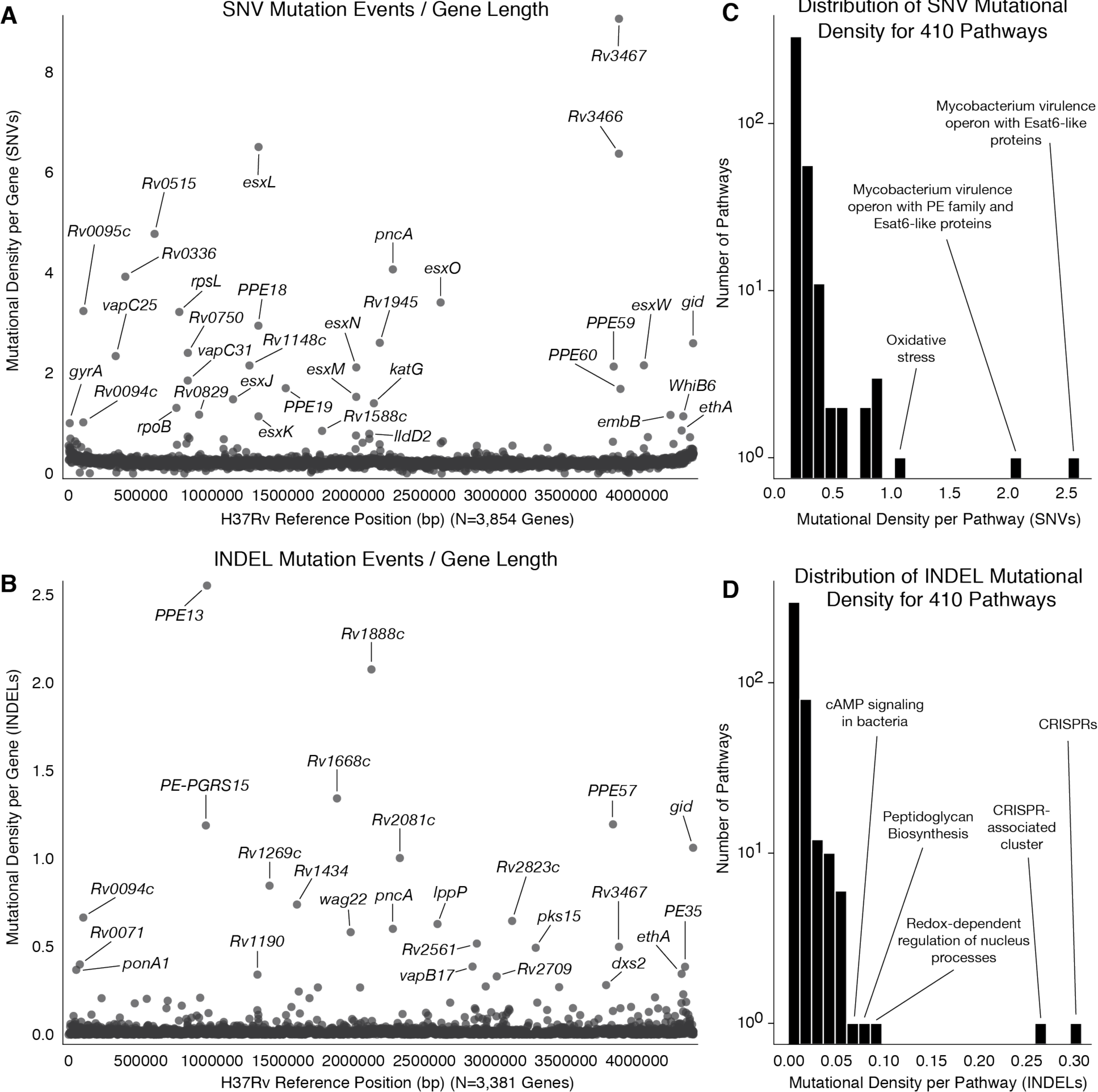
SNV and INDEL mutational density per gene. (**A**) The homoplasy scores for all SNVs within each gene were aggregated to approximate all SNV mutation events (independent arisals) that occurred within the gene body then normalized by the gene length (**Materials and Methods**). **Table S8** contains the calculations for each gene as well as columns for *# SNVs*, *Synonymous Homoplasy Score*, and *Non-Synonymous Homoplasy Score*. (**B**) A similar computation was carried out for INDELs in which homoplasy scores for all INDELs within each gene were aggregated and normalized by gene length (**Materials and Methods**). **Table S9** contains the calculations for each gene as well as *# INDELs*, *Inframe Homoplasy Score*, and *Frameshift Homoplasy Score*. (**C**) Homoplasy scores for all SNVs were aggregated at the level of pathways then normalized by the gene lengths for each gene set (**Table S10**, **Materials and Methods**). (**D**) Homoplasy scores for all INDELs were aggregated at the level of pathways then normalized by the gene lengths for each gene set (**Table S11**, **Materials and Methods**).

Among the calculations for SNVs (**Figure 5A**, **Table S8**), several outlier genes are involved in the acquisition of antibiotic resistance (*gyrA*, *rpoB*, *rpsL*, *gid*, *katG*, *pncA*, *embB*) (Farhat et al., 2013; Manson et al., 2017). Additionally, several outliers belonged to the ESX protein family (*esxL*, *esxO*, *esxN*, *esxM*, *esxW*) which are involved in host-pathogen interactions (Uplekar et al., 2011) and the PE/PPE protein family (*PPE18*, *PPE19*, *PPE59*, *PPE60*) which include antigenic proteins (Brennan, 2017). For INDELs (**Figure 5B**, **Table S9**), outliers included the antibiotic resistance loci: *pncA*, *gid* (Coll et al., 2018; Manson et al., 2017) and additional members of the PE/PPE family (*PPE13*, *PE-PGRS15*, *PPE57*). Next, we extended this analysis for SNVs & INDELs at the pathway level by aggregating Hs across different gene sets belonging to 410 pathways (**Materials and Methods**). The pathway with the most mutational density per SNVs belonged to a *Mycobacterium* virulence operon with Esat6-like proteins (**Figure 5C**, **Table S10**), while the pathway most enriched for mutational density per INDELs belonged to the CRISPR associated cluster that contains the aforementioned putative antigen Rv2823c (**Figure 5D**, **Table S11**).

## DISCUSSION

As MTBC evolved into a professional pathogen from a saprophytic mycobacterium, it underwent step-wise adaptation to the intracellular environment. This adaptation is thought to comprise genome contraction, expansion of specific gene families especially toxin-antitoxin systems, the type VII secretion systems, and the PE-PPE gene family, as well as gene modification through mutation (Gagneux, 2018). Population genetic studies of MTBC have largely concluded that the modern MTBC genome is under purifying selection with most newly fixed diversity attributable to antibiotic selection pressure (Brynildsrud et al., 2018; Chiner-Oms et al., 2019; Holt et al., 2018; Vargas et al., 2021). It has thus been suggested that MTBC has reached a pathogenic fitness peak (Pepperell et al., 2013). Here, we update this view by analyzing the largest to date collection of MTBC genome sequences characterizing the timing and pattern of genetic variation acquisition across the phylogeny. We find 4,980 SNVs, 993 non-SSR related INDELs, and 45 HT regions to have evolved in a parallel manner with high frequencies suggestive of an adaptive role. Although a subset of this variation can be linked to resistance based on known genetic determinants, the majority has no known association with resistance. Among the highest scoring variants we find proteins that encode putative antigens (*esxL*, *esxW, Rv2823c*) (Tak et al., 2021), other PE/PPE proteins (*PPE54* and *PPE18*) (Vargas et al., 2021), toxin-antitoxin bicistrons (*vapC2*, *mazF6*) and ESX-1 system (*espK, espA, espR*) strongly suggestive of a role in virulence (Garces et al., 2010). The highest scoring variants also heavily overrepresent intergenic regions (20%, 22%, and 26% of putatively adaptive SNVs, non-SSM INDELs, and HTs respectively) even though intergenic regions constitute only 10% of the genome by length. Putatively adaptive transcriptional variants appear to converge with protein variants in impacting ESX-1 function. We identify a substantial proportion of putatively adaptive variation to be acquired recently and on par with acquisition of resistance related variants, suggesting that modern MTBC continues to refine its virulence strategies likely in the context of a dynamic host environment.

Phase variation was recently recognized to mediate MTBC drug-tolerance through frameshifts in the glycerol kinase gene *glpK* that likely act by altering the metabolic state of the cell (Bellerose et al., 2019; Safi et al., 2019). In other bacterial pathogens, phase variation can alter antibiotic efficacy and the immunogenicity of cell surface proteins through altered transcription, translation and/or the creation of protein diversity (Van Der Woude and Bäumler, 2004). Here, we take a genome-wide approach to assess the frequency and impact of phase variation in MTBC. We measure the frequency of INDEL acquisition in HTs at 38x the rate observed for SNVs in clinical isolates. Based on *in vitro* measurements, we estimate the frameshift rate under expected neutral conditions at 1.1 × 10^−5^ frameshifts/HT/year, ∼100x the rate previously reported MTBC SNV acquisitions (Walker et al., 2013). The discrepancy between the *in vitro* and observed event rate in HTs in clinical isolates is likely attributable to INDEL reversions. Remarkably despite the undercounting of INDEL events in HTs, more than 12% of all INDEL events observed in the MTBC clinical isolate phylogeny occur in an HT region. We find a few examples of frequent SSM in non-HT SSR regions, *e.g.*, in *ponA1,* a gene previously identified to modulate growth in the presence of the drug rifampicin (Farhat et al., 2013). However, we measure a substantially lower rate of INDELs in the latter regions compared with HTs (**Figure 1B**, **Figure S7**). Using a GWAS approach, we discover a subset of frameshifts in HTs to be associated with antibiotic resistance. These include genes of unknown function Rv3413c and Rv2264c as well as an HT upstream of lysyl-tRNA synthetase *lysX*. This gene is conditionally essential for bacterial growth *in vivo*, its higher expression correlates positively with virulence in clinical isolates, and in *M. avium hominis lysX* mutants associate with resistance to cationic antimicrobials and increased inflammatory response after macrophage infection (Kirubakar et al., 2020; Montoya-Rosales et al., 2017; Sassetti and Rubin, 2003). Hence the frameshifts in the HT upstream of *lysX* may plausibly affect both antibiotic resistance and virulence in MTBC.

Multiple different pressures may differentially select for variants related to ESX-1 activity. This secretion system influences virulence and antigenicity in MTBC (Garces et al., 2010; Lim et al., 2022) by controlling the secretion of the immunodominant antigens ESAT-6 (*esxA*) and CFP-10 (*esxB*) (Covert et al., 2001; Guinn et al., 2004; Hsu et al., 2003), stimulating the innate immune response and cytokine secretion (Pandey et al., 2009, p. 2; Stanley et al., 2007), and promoting the intracellular growth of the pathogen (Lewis et al., 2003; Stanley et al., 2003). Through modulating the immune response, as well as cellular permeability (Garces et al., 2010), ESX-1 function may also influence antibiotic activity or resistance (Torres Ortiz et al., 2021). Indeed, we identified phase variants that truncate *espK*, an ESX-1 associated gene that when disrupted *in vitro* promotes bacterial growth (DeJesus et al., 2017) to associate with resistance. In contrast, INDELs that reduce the expression of the *espACD* operon were not associated with the resistant phenotype, suggesting that another host-derived pressure may be responsible for selecting these variants. These indels might be expected to reduce bacterial fitness, as deletion of *espA* abrogates secretion of ESAT-6 and CFP-10 and attenuates growth in mice to a similar degree as deletion of the ESX-1 locus (Fortune et al., 2005). However, lower levels of ESX-1 function could also result in reduced antigen presentation and/or cytokine production, thus aiding immune evasion (Clemmensen et al., 2017). We thus hypothesize that multiple modes of phase variation tune ESX-1 activity to optimize growth, survival, or transmission. These states may influence antibiotic susceptibility through modulation of growth and membrane permeability, or by altering the local environment. These hypotheses are testable in *in vivo* experimental systems.

This analysis is not without limitations. First is our inability to functionally validate all novel associations due to the time and resources needed to manipulate Mtb genetically *in vitro*. Instead, we provide a proof of concept validation of transcriptional regulation for one HT candidate in the transcriptional start site of *espA*. Second is our inability to assess adaptive INDELs in non-HT SSR regions as they vary in their sequence composition and the expected rate of SSM, thus challenging our ability to simulate neutral evolution in these regions. Similarly, it is difficult to account for the reversibility of INDELs in SSR regions, and it is possible that some homoplasic variants represent a combination of mutation and reversion, as opposed to two distinct arisals. Regardless, the reported Hs values still represent the number of independent mutational events observable at a site. In this work, we also make the assumption that SNV mutation rates are homogeneous outside of SSR regions. We recognize that many forces likely determine the neutral mutation rate across the genome including GC content, repetitive sequence, and transcription coupled repair to name a few factors. Driving both extremes of evolutionary rates are forces of positive and purifying selection respectively that shape the genome. The approach we take in simulating neutral evolution is only a useful approximation to gauge the very extreme rates of evolution. It is likely that regions with seemingly borderline rates of Hs may also have functional consequences, and at the other extreme are genes under purifying selection that are beyond the scope of this work.

In summary, in this work we present evidence that MTBC genomes are strongly and regionally shaped by positive selection not only to modulate the resistance phenotype but likely also virulence mechanisms. We hypothesize that phase variation in ESX-1 system of MTBC can act as a toggle between antigenicity and survival in the host. The ongoing regional evolution of MTBC suggests that the host environment in MTBC infection is dynamic, including potentially opposing forces that shape transmissibility and survival in host. Overall the insights gained in this analysis can inform vaccine design and host and pathogen-directed therapy against MTBC that have recently been expanded to include ESX-1 targeting compounds (Cole, 2016).

## MATERIALS AND METHODS

### Sequence Data

We initially downloaded raw Illumina sequence data for 33,873 clinical isolates from NCBI (Benson et al., 2000). We identified the BioSample for each isolate and downloaded all of the associated Illumina sequencing runs. Isolates had to meet the following quality control measures for inclusion in our study: (i) at least 90% of the reads had to be taxonomically classified as belonging to MTBC after running the trimmed FASTQ files through Kraken (Wood and Salzberg, 2014) and (ii) at least 95% of bases had to have coverage of at least 10x after mapping the processed reads to the H37Rv reference genome (Genbank accession: NC_000962).

### Illumina Sequencing FastQ Processing and Mapping to H37Rv

The raw sequence reads from all sequenced isolates were trimmed with version 0.20.4 Prinseq (settings: -min_qual_mean 20) (Schmieder and Edwards, 2011) and then aligned to H37Rv with version 0.7.15 of the BWA mem algorithm using the -M settings (Li and Durbin, 2009). The resulting SAM files were then sorted (settings: SORT_ORDER = coordinate), converted to BAM format, and processed for duplicate removal with version 2.8.0 of Picard (http://broadinstitute.github.io/picard/) (settings: REMOVE_DUPLICATES = true, ASSUME_SORT_ORDER = coordinate). The processed BAM files were then indexed with Samtools (Li et al., 2009). We used Pilon (settings: --variant) on the resulting BAM files to generate VCF files that contained calls for all reference positions corresponding to H37Rv from pileup (Walker et al., 2014).

### Empirical Score for Difficult-to-Call Regions

We assessed the congruence in variant calls between short-read Illumina data and long-read PacBio data for a set of isolates that underwent sequencing with both technologies (Marin et al., 2022). Using 31 isolates for which both Illumina and a complete PacBio assembly were available, we evaluated the empirical base-pair recall (EBR) of all base-pair positions of the H37rv reference genome. For each sample, the alignments of each high confidence genome assembly to the H37Rv genome were used to infer the true nucleotide identity of each base pair position. To calculate the empirical base-pair recall, we calculated what percentage of the time our Illumina based variant calling pipeline, across 31 samples, confidently called the true nucleotide identity at a given genomic position. If Pilon variant calls did not produce a confident base call (*Pass*) for the position, it did not count as a correct base call. This yields a metric ranging from 0.0–1.0 for the consistency by which each base-pair is both confidently and correctly sequenced by our Illumina WGS based variant calling pipeline for each position on the H37Rv reference genome. An H37Rv position with an EBR score of x% indicates that the base calls made from Illumina sequencing and mapping to H37Rv agreed with the base calls made from the PacBio *de novo* assemblies in x% of the Illumina-PacBio pairs. We masked difficult-to-call regions by dropping H37Rv positions with an EBR score below 0.9 (or 90%) as part of our variant calling procedure. Full details on the data and methodology can be found elsewhere (Vargas et al., 2021).

### Variant Calling

#### SNP Calling

To prune out low-quality base calls that may have arisen due to sequencing or mapping error, we dropped any base calls that did not meet any of the following criteria: (i) the call was flagged as *Pass* by Pilon, (ii) the mean base quality at the locus was >20, (iii) the mean mapping quality at the locus was >30, (iv) none of the reads aligning to the locus supported an insertion/deletion (indel), (v) a minimum coverage of 20 reads at the position, and (vi) at least 75% of the reads aligning to that position supported 1 allele (using the *INFO.QP* field which gives the proportion of reads supporting each base weighted by the base and mapping quality of the reads, *BQ* and *MQ* respectively, at the specific position). A base call that did not meet all filters (i) – (vi) was inferred to be low-quality/missing (**Figure S2**).

#### INDEL Calling

To prune out low-quality INDEL variant calls, we dropped any INDEL that did not meet any of the following criteria: (i) the call was flagged as *Pass* by Pilon, (ii) the maximum length of the variant was 10bp, (iii) the mean mapping quality at the locus was >30, (iv) a minimum coverage of 20 reads at the position, and (v) at least 75% of the reads aligning to that position supported the INDEL allele (determined by calculating the proportion of total reads *TD* aligning to that position that supported the insertion or deletion, *IC* and *DC* respectively). A variant call that met filters (i), (iii), and (iv) but not (ii) or (v) was inferred as a high-quality call that did not support the INDEL allele. Any variant call that did not meet all filters (i), (iii), and (iv) was inferred as low-quality/missing.

### Lineage Typing and Classifying Isolates into Groups

After excluding 1663/33873 isolates that had missing calls > 10% SNP sites , we determined the global lineage of each isolate (*N* = 32210) using base calls from Pilon-generated VCF files and a 95-SNP lineage-defining diagnostic barcode (**Figure S2**) (Freschi et al., 2021). We further excluded 290 isolates that had no lineage call or more than one lineage call (low-quality calls at lineage-defining SNP sites or a rare SNP call characterized as monophyletic for another lineage in the SNP barcode), and 35 isolates that had L7 lineage calls (**Figure S2**). Our remaining 31885 isolates were typed as: L1 (2815), L2 (8090), L3 (3398), L4 (17388), L5 (98), L6 (96). We aimed to cluster isolates into groups of no more than 8,000 isolates based on lineage & sub-lineage to achieve feasible phylogeny construction runtimes so we further divided L4 isolates based on sub- lineage calls. We excluded 457 isolates that were typed as L4 but did not have any sub-lineage calls. We analyzed the sub-lineage calls of the remaining 16931 L4 isolates and grouped isolates according to sub-lineages that were located next to each other on the L4 phylogeny (Freschi et al., 2021). We grouped the L4 isolates into three groups: L4A (sub-lineages 4.1.x & 4.2.2.x, *N* = 5839), L4B (sub-lineage 4.2.1.2.x, *N* = 6958), and L4C (sub-lineage 4.2.1.1.x, *N* = 4134) where *.x* is a place-holder for any further resolution on the sub-lineage call under the hierarchical lineage typing scheme (Freschi et al., 2021).

### SNP Genotypes Matrix

A schematic diagram outlining the following steps is given in **Figure S2**. First, we detected SNP sites at 899,035 H37Rv reference positions (of which 64,950 SNPs were not biallelic) among our global sample of 33,873 isolates. We constructed a 899,035x33,873 genotypes matrix (coded as 0:A, 1:C, 2:G, 3:T, 9:Missing) and filled in the matrix for the allele supported at each SNP site (row) for each isolate, according to the *SNP Calling* filters outlined above. If a base call at a specific reference position for an isolate did not meet the filter criteria that allele was coded as *Missing*. We excluded 20,360 SNP sites that had an EBR score <0.90, another 9,137 SNP sites located within mobile genetic element regions (e.g. transposases, intergrases, phages, or insertion sequences) (Comas et al., 2010; Vargas et al., 2021), then 31,215 SNP sites with missing calls in >10% of isolates, and 2,344 SNP sites located in overlapping genes (coding sequences). These filtering steps yielded a genotypes matrix with dimensions 835,979x33,873. Next, we excluded 1,663 isolates with missing calls in >10% of SNP sites yielding a genotypes matrix with dimensions 835,979x32,210 (Coll et al., 2018). We used an expanded 96-SNP barcode to type the global lineage of each isolate in our sample (Freschi et al., 2021). We further excluded 325 isolates that either did not get assigned a global lineage, assigned to more than one global lineage, or were typed as lineage 7. We then excluded 41,760 SNP sites from the filtered genotypes matrix in which the minor allele count = 0 which resulted in a 794,219x31,885 matrix. To provide further MTBC lineage resolution on the lineage 4 isolates, we required an MTBC sub-lineage call for each lineage 4 isolate. We excluded 457 isolates typed as global lineage 4 but had no further sub-lineage calls and then again excluded 11,654 SNP sites from the filtered genotypes matrix in which the minor allele count=0. The genotypes matrix used for downstream analysis had dimensions 782,565x31,428, representing 782,565 SNP sites across 31,428 isolates (**Figure S2**). The global lineage (L) breakdown of the 31,428 isolates was: L1=2,815, L2=8,090, L3=3,398, L4=16,931, L5=98, L6=96.

### INDEL Genotypes Matrix

We detected 53,167 unique INDEL variants within 50,576 H37Rv reference positions among our global sample of 33,873 isolates. We constructed a 53,167x33,873 genotypes matrix (coded as 1:high quality call for the INDEL allele, 0:high quality call not for the INDEL allele, 9:Missing) and filled in the matrix according to whether the INDEL allele was supported for each INDEL variant (row) for each isolate, according to the *INDEL Calling* filters outlined above. If a variant call at the reference position for an INDEL variant did not meet the filter criteria that call was coded as *Missing*. We excluded 2,006 INDELs that had an EBR score <0.90, another 694 INDELs located within mobile genetic element regions, then 207 INDELs located in overlapping genes (coding sequences). These filtering steps yielded a genotypes matrix with dimensions 50,260x33,873. Next, we excluded any isolate that was dropped while constructing the SNP genotypes matrix to retain the same 31,428 isolates as described above. Finally, we excluded 2,835 INDELs in which the alternate allele count=0. The genotypes matrix used for downstream analysis had dimensions 47,425x31,428 (**Figure S1A**).

### Phylogeny Construction

To generate the phylogenies, we first merged the VCF files of the isolates in each group (L1, L2, L3, L4A, L4B, L4C, L5, L6) with bcftools (Li et al., 2009). We then removed repetitive, antibiotic resistance and low coverage regions (Freschi et al., 2021). We generated a multi-sequence FASTA alignment from the merged VCF file with vcf2phylip (version 1.5, https://doi.org/10.5281/zenodo.1257057). We constructed the phylogenetic trees with IQ-TREE (Nguyen et al., 2015). For all groups, we used the *mset* option to restrict model selection to GTR models (`-mset GTR`), and specified 1000 bootstrap replicates for both ultrafast bootstrap and SH-aLRT algorithms to compute support values (`-bb 1000 -alrt 1000`). To construct phylogenies for groups L1, L2, L3, L4A, L4B & L4C, we specified the substitution model as GTR+F+I+R (`-m GTR+F+I+R`). To construct phylogenies for groups L5 & L6, we implemented the automatic model selection with ModelFinder Plus (`-m MFP`) (Kalyaanamoorthy et al., 2017). The runtimes to construct the phylogenies were: L1 (2 days, 1.5 hours), L2 (63 days, 9 hours), L3 (11 days, 20 hours), L4A (6 days, 11 hours), L4B (6 days, 18 hours), L4C (2 days, 18 hours), L5 (4 minutes), L6 (2.5 minutes). Upon closer inspection of the phylogenies, we observed that a handful of isolates (14/31428) were misclassified based on the SNP barcode. The misclassified isolates belonged to the following groups: L1 (3), L2 (4), L3 (2), L4A (1), L4B (0), L4C (4), L5 (o), L6 (0). The small number of mistyped isolates did not affect our inferences so we kept these phylogenies for downstream analyses.

### Assessment of Parallel Evolution for SNVs

To quantify the number of independent arisals for each SNV, we used the SNP genotypes matrix in conjunction with the phylogenies for each isolate group (**Figure S1B**). We used an ancestral reconstruction approach to quantify the number of times each SNV arose independently within each phylogeny using SNPPar (**Figure S5B**) with options: `–-sorting intermediate –- no_all_calls ----no_homoplasic` (Edwards et al., 2020). We parsed the SNPPar output files `all_muation_events.tsv` and `node_sequences.fasta` to check each mutation reported in the mutation events table against the inferred sequences at the nodes of the phylogeny and the isolates sequences. Mutations that were not found in the sequences were discarded, the number of reported mutation events not located between inferred node/isolate sequences is broken down by phylogeny as follows: L1 (447), L2 (2472), L3 (392), L4A (839), L4B (1177), L4C (559), L5 (2), L6 (3). We then parsed the filtered *mutation events* tables corresponding to each isolate group and counted the number of times each unique SNV in our dataset was inferred to have arisen, counting only the number of times that the major allele (ancestor call) mutated toward the minor allele (derived call) for each SNV (**Figure S5B**). This yielded a *homoplasy score* or an estimate for the number of independent arisals for each SNV across all 31,428 isolates (**Table S1**, **Table S2**). We note that 1,920/836,901 SNVs in our SNP genotypes matrix had a *homoplasy score* = 0, this was likely due error in the ancestral reconstructions, or may have been the result of sub-setting isolates into groups before running ancestral reconstruction (i.e. if an SNV is fixed in isolates belonging to one of the phylogenies but not called in any other isolates, no mutation event would be reported). These SNVs were dropped from downstream analysis.

### Assessment of Parallel Evolution for INDELs

To quantify the number of independent arisals for each INDEL, we developed a simple method to count the number of times each a given allele “breaks” the phylogenies (**Figure S5C**). If a given minor/alternate allele is observed in two separate parts of a phylogeny, then we can assume that this allele arose twice in pool of isolates used to construct the tree. If the minor/alternate allele is observed in three separate parts of the phylogeny, then we assume that the allele arose independently three times. We extended this idea to count the total number of times a given minor/alternate allele arises within a phylogeny. To do this we specify a minor/alternate allele of interest and code the phylogeny tips (according to weather the corresponding isolates harbor the allele) as follows: minor/alternate allele = 1, major/reference allele = 0, low quality call = 9. We create a vector from the coded phylogeny tips and then count the number of times each consecutive string of 1’s appears in the vector. These consecutive 1’s (“1 blocks”) must be separated by 0’s on either side, and the number of 0’s required in between the strings of 1’s is controlled by the *spacer* parameter. If spacer = 1, then only one 0 is required in between 1 blocks to count different arisals. If spacer = 2, then two 0’s are required between 1 blocks to count them as separate arisals (**Figure S5C**). We allowed the presence of 9’s in the 1 blocks as long as a 1 was present in the block. As an example, suppose a phylogeny of 15 isolates had tips coded as [0,0,1,1,0,1,0,0,0,1,1,1,0,0,0] for a given allele. If spacer = 1, then [0,0,**1**,**1**,0,**1**,0,0,0,**1**,**1**,**1**,0,0,0] would correspond to three 1 blocks and we would infer three independent arisals or a *homoplasy score* = 3. If spacer = 2, then [0,0,**1**,**1**,**0**,**1**,0,0,0,**1**,**1**,**1**,0,0,0] would correspond to two 1 blocks and we would infer two independent arisals or a *homoplasy score* = 2. Higher values of the spacer parameter yield more conservative estimates for *homoplasy score* calculations.

We calculated a *homoplasy score* by counting these topology disruptions (TopDis) or “blocks” for SNVs using the SNP genotypes matrix in conjunction with the phylogenies for each isolate group to assess the number of independent arisals for each mutation observed, coding the tips as 1 if they carried the minor allele for each SNV (**Figure S5C**, **Figure S1C**). We computed these *homoplasy scores* for different values of the spacer parameter (1-6) to assess the congruence of these estimates with the *homoplasy scores* computed from the ancestral reconstructions (**Figure S6**). The results were concordant between both methods, although TopDis appeared to overestimated the *homoplasy score* for some SNVs with spacer = 1 and spacer = 2 (**Figure S6A-B**). These results validated our approach for computer *homoplasy scores* using TopDis. To compute the *homoplasy scores* for INDELs, we conservatively chose spacer = 4 at which point the *homoplasy score* for each SNV computed from TopDis appeared to be equal or less than the *homoplasy score* computed from SNPPar (**Figure S6D**). To quantify the number of independent arisals for each INDEL, we used the INDEL genotypes matrix in conjunction with the phylogenies for each isolate group as input to TopDis with spacer = 4 (**Figure S1D**), coding the tips as 1 if they carried the alternate allele for each INDEL (**Figure S5C**). We note that 1,119/47,425 indels had *homoplasy score* = 0, this may have been the result of sub-setting isolates into groups before running TopDis (i.e. if an INDEL is fixed in isolates belonging to one of the phylogenies but not called in any other isolates, no “block” would be observed) or if the INDEL alternate allele was only present at the ends of the coded phylogeny tips vector. These INDELs were dropped from downstream analysis.

### Homoplasy Simulations for SNVs & INDELs

We aimed to assess the frequency with which a specific mutation would repeatedly arise by chance given the phylogenies used to related the isolates in our dataset. We assumed a constant population size model which has previously been used to estimate the molecular clock rate of Mtb (Menardo et al., 2019). Menardo et al. estimated the molecular clock rate for Mtb using a Bayesian phylogenetic approach under two different coalescent priors, (1) constant population size and (2) exponential population growth, for 21 datasets of Mtb strains that showed stronger temporal signal than expected by chance by preforming a date randomization test on the corresponding phylogenies (Menardo et al., 2019). They found that 14/21 datasets rejected the constant population size model, however the results were only moderately influenced by the tree prior and their molecular clock estimates were robust different demographic models.

We converted the phylogeny branches to time and assumed that neutral mutations arise on the genome according to a Poisson distribution. To simulate the expected number of arisals (Hs) for neutral point mutations in our dataset, we simulated mutations on the branches of the eight phylogenies that relate all of the isolates in our sample. First, we extracted the branch lengths (*b*) from each tree along with the length of the SNP concatenate (*l*) used to construct each tree *s*. Then, for each branch *i* for each tree *s*: (1) we drew a molecular clock rate μ_’_∼*U*(0.3,0.6) (assuming a neutral rate of 0.5 SNPs/genome/year (Vargas et al., 2021; Walker et al., 2013), (2) we converted the branch length to years *t*_’_ = (*b*_’_ × *l*_(_)/μ_’_, (3) we assumed neutral point mutations accrued according to the molecular clock and drew the rate according to *v*_*i*_∼*U*(0.3,0.6), (4) we assumed that neutral mutations on the genome follow a Poisson distribution and calculated λ for each branch as λ_’_ = *t*_’_ × *v*_*i*_, (5) we drew the number of mutations expected to occur on *b*_’_ as *n*_*i*_∼*Poiis*(λ_*i*_), (6) assuming that neutral mutations occur anywhere along the 4Mbp genome of Mtb with equal likelihood, we randomly chose *n*_*i*_ positions between 1-4,000,000 to simulate the positions where each mutation occurred.

We repeated this process for all branches across all trees and kept track of the number of times each position between 1-4,000,000 was selected to approximate Hs for each position. This resulted in an approximate Hs for each position that was selected at least once (number of times each position was selected) and a distribution of the number of positions for increasing values of Hs. Lastly, we repeated all of the steps above 100 times to get the probability that a neutral mutation arises at a specific position ≥ *Hs* by taking the median (for each *Hs*) across 100 simulations. This then gave us a proportion of genome positions that were homoplastic by chance (*Hs* ≥ 2). By taking the median across 100 simulations for increasing values of Hs, we observed that *P*(*Hs* ≥ 5) < 0.002 and used *Hs* = 5 as a threshold for assessing which SNVs were unlikely to have arisen repeatedly by chance alone. As neutral insertions and deletions generated by non-SSM mechanisms are expected to occur more rarely than SNVs, we conservatively used this threshold to further analyze INDELs in non-HT and non-SSR genomic regions that were unlikely to have repeatedly arisen by chance.

We modified the process above to simulate neutral mutational density for each gene. For each gene of length *l*, we normalized the mutation rate *v*_*i*_ to account for gene length by multiplying it by *l*_*i*_/*L* where L is the length of the genome (4Mbp). Then we simulated mutations that occur on each branch across all phylogenies and added each mutation to a count to calculate number of mutations (independent arisals) that occurred in *g*_*i*_ under neutrality. We repeat this process 100 times for each gene *g*_*i*_ to get an average number of neutral mutations that arise in each gene *g*_*i*_ across 100 simulations and divide by the gene length *l*_*i*_ to get the neutral mutational density for *l*_*i*_. The neutral mutation densities across all genes ranged from 0.42-0.45, the mean mutational density was 0.44, and the 99.8^th^ percentile of mutational densities across genes was 0.449.

### Media

*Mycobacterium tuberculosis* H37Rv and *Mycobacterium smegmatis* were both grown in 7H9 broth with 0.05% Tween 80, 0.2% glycerol, and OADC (oleic acid-albumin-dextrose-catalase; Becton, Dickinson); transformants were selected on 7H10 plates with 0.5% glycerol and OADC. When needed, the following supplements were added: kanamycin (25 μg/ml), hygromycin (50 μg/ml), anhydrotetracycline (aTc).

### Recombineering Single Nucleotide espA Mutant

Mtb harboring pKM402 (Ioerger et al., 2013) and pKM427 (Murphy, 2021) were grown in 30 ml 7H9 media containing OADC, 0.2% glycerol, 0.05% Tween 80, and 25 μg/ml kanamycin. Atc was added to a final concentration of 500 ng/ml at OD ∼0.4, Electroporation was performed as described (Murphy, 2021) using 2 ug espA target oligo (GGCCTACAGTCTGGCTGTCATGCTTGGCCGATGTCAACAGTTTTTTCATGCTAAGCAGATCGTCAGTTTTGAGTTCGTGAAGACGG) and 200 ng hygR repair oligo (CGGTCCAGCAGCCGGGGCGAGAGGTAGCCCCACCCGCGGTGGTCCTCGACGGTCGCCGCG). Candidate clones were expanded in into 4 mL 7H9-OADC-Tween with 50 ug/mL hygromycin. The upstream region of Rv3616 was amplified by PCR using the following primers: GACCGGGATGTAGGTCAGGTC) and GCTAGGTGTTTAGCGGACGCG. The PCR product was sequenced with GCTAGGTGTTTAGCGGACGCG as a primer to confirm the presence of the mutation.

### RNA Extraction

10 mL of WT and mutant H37Rv were grown at 37°C in 7H9-OADC-Tween to an OD ∼0.6. Immediately prior to harvesting RNA, 40 mL of guanidine isothiocyanate buffer was added to the culture (5 M guanidine isothiocyanate, 0.5 % N-lauryl sarcosine, 25mM Tri-Sodium citrate, 0.1 M beta -mercaptoethanol, 0.5% Tween, pH = 7.0). Bacteria were collected by centrifugation at 4,000 rpm for 10 minutes at 4°C, resuspended in 500 uL of Trizol reagent, and lysed bylysing matrix B (MP Bio) and bead beating three times at 6.5 M/s for 45 seconds. After centrifugation, 100 uL of chloroform was added to the supernatant, inverted several times, and incubated at room temperature for 3 minutes. Samples were then centrifuged at 10,000 x g at 4°C for 15 minutes. RNA was then extracted using Zymo Research Direct-zol RNA Miniprep Plus Kit. Samples were processed according to manufacturer’s instructions, including 15 minute on-column DNase digestion. After eluting in 50 uL, an additional DNase digestion was performed using NEB RNase-free DNase I, bringing the total volume of the reaction up to 100 uL. Samples were incubated at 37°C for 2 hours. 100 uL of the reaction was then added to 400 uL of Trizol reagent, to which 500 uL of ethanol was then added. RNA extraction with Direct-zol kit was repeated as before, but this time skipping the on-column DNase digestion. Samples were eluted in 50 uL of water and the concentration of each sample was determined via NanoDrop.

The RNA extracted from the *espA* mutant was sequenced on an Illumina 4000 in paired-end mode, with a read-length of 2x150 bp. Two runs were performed, and 3 replicates each for the *espA* mutant (+1 bp insertion in homopolymer) and wild-type (*M. tuberculosis* H37Rv) were collected on each run. The reads were mapped to the H37Rv genome (Genbank accession NC_000962.2) using BWA (v0.7.12), and read counts for each ORF (open reading frame) were tabulated. The R package DeSeq (Love et al., 2014) was used to analyze the counts and identify differentially expressed genes as genes with an adjusted P-value < 0.05 (after multiple-tests correction). DeSeq internally normalizes the count data by computing scaling factors for each dataset. The model was fit with 2 covariates, strain and run, and the statistical analysis was based on the strain coefficient (as contrast), to evaluate the average effect of the *espA* mutant on the counts for each gene relative to the WT samples from the same run.

### RNAseq Library Preparation

250ng of total RNA was processed using the Illumina Ribo-Zero Plus rRNA Depletion Kit and NEBNext® Ultra II Directional RNA Library Prep Kit for Illumina. Adaptor ligated DNA was PCR enriched for 9 cycles according to the protocol using indexed primers from NEBNext Multiplex Oligos for Illumina. Samples were purified using SPRIselect Beads at each clean-up step. Prepared libraries were diluted to equal concentrations and pooled at a concentration of 30 nM. Samples were processed on an Illumina HiSeq 4000 machine with a 2 x 150 basepair sequencing configuration.

### Plasmid Construction

A homopolymer frameshifting reporter was constructed from a hygromycin resistant pDE43- MCtH vector, which is a version of pDE43-MCK with a swapped antibiotic marker (Addgene plasmid #49523; (Kim et al., 2013)). Using Gibson assembly, the homopolymer sequence from *glpK* (Rv3696c) along with 79 basepairs of flanking sequence (40 basepairs preceding, and 39 basepairs following, the homopolymer) was fused to an out-of-frame kanamycin resistance cassette, such that the addition of a single nucleotide insertion would produce an in-frame kanamycin resistance gene, all of which is driven by a P16 mycobacterial-specific promoter (a gift from Dirk Schnappinger).

### Fluctuation Analysis

*Mycobacterium smegmatis* harboring the homopolymer-frameshifting reporter was thawed from glycerol stock and grown in 4 mL 7H9-OADC-Tween to an OD ∼1.0. This culture was split and diluted into 20 parallel cultures, each with an OD = 0.01. These cultures were rotated at 37°C for ∼20 hours. Total bacterial numbers were determined by plating on 7H10 plates with OADC and 0.5% glycerol. To enumerate frameshifted mutants, entirety of each culture was plated on 7H10 with 25 ug/mL kanamycin. Plates were incubated at 37°C for 4-5 days prior to counting colonies. In a subset of kanamycin resistant colonies, frameshifts were verified by PCR with the following primers: GCTCGAATTCACTGGCCATGCATC) and GATCCTGGTATCGGTCTGCGATTC.

The PCR product was then sequenced using GATCCTGGTATCGGTCTGCGATTC as a primer. After accounting for the proportion of kanamycin resistant colonies that contained a frameshifted homopolymer (20/28), the mutation rate was calculated as described by (Gillet-Markowska et al., 2015, p.).

### Homoplasy Simulations for INDELs in a Homopolymer Tract

Similar to the simulations for point mutations, we assumed that frameshift mutations arise within the HT according to a Poisson distribution and assign frameshift mutations (insertions and deletions) to the branches of our phylogenies by drawing from a Poisson distribution with lambda modified by the length of each branch and the experimentally derived mutation rate for frameshifts within an HT (**Figure S8**). To simulate the number of expected number of arisals (Hs) for frameshift mutations (FS) within homopolymeric tracts (HT), we simulated frameshifts for a 7bp HT on the branches of the eight phylogenies that related all of the isolates in our sample. We calculated a frameshift rate for neutral frameshifts in a HT using the lower and upper bound mutation rates reported from the *Mycobacterium smegmatis* fluctuation analysis 2.18 × 10^−8^ and

4.09 × 10^−8^ mutation rates/cell/division (**Materials and Methods**). We note that Mtb doubles once every 24 hours in liquid culture (Gill et al., 2009). We added stochasticity to the doubling time by incorporating a term 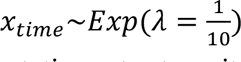 and calculated doublings per day as 24/(24 + *x_time_*). We converted these mutation rates to units of FS/HT/year as follows:

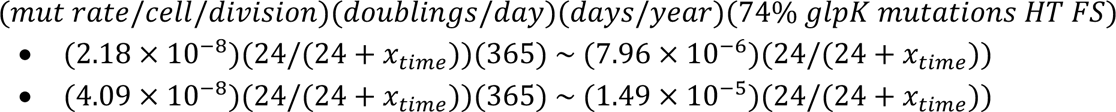

to get the lower bound (*FS*_*lower*_) and upper bound (*FS*_*upper*_) FS/HT/year neutral rates, respectively.

As with simulations for point mutations, we extracted all of the branch lengths (*b*) from each tree along with the length of the SNP concatenate (*l*) used to construct each tree *s*. Then, for each branch *i* for each tree *s*: (1) we drew a molecular clock rate μ_*i*_∼*U*(0.3,0.6) (assuming a neutral rate of 0.5 SNPs/genome/year (Vargas et al., 2021; Walker et al., 2013), (2) we converted the branch length to years *t*_*i*_ = (*b*_*i*_ × *l*_(_)/μ_*i*_, (3) we drew *x*_*time*_∼*Exp*(λ = 1/10) and calculated *FS*_*lower*_ and *FS*_*upper*_, (4) we drew a FS neutral mutation rate according to *v*_*i*_∼*U*(*FS*_*lower*_, *FS*_*upper*_), (5) we assumed that neutral FS in the HT follow a Poisson distribution and calculated λ for each branch as λ_*i*_ = *t*_*i*_ × *v*_*i*_, (6) we drew the number of FS expected to occur within HT on *b*_*i*_ as *n*_*i*_∼*Pois*(λ_*i*_), (7) we ran a Bernoulli trial with *p* = 0.5 to assign *n*_*i*_ as an insertion (+) or deletion (-) on *b*_*i*_ (**Figure S8**) and assigned an INDEL genotype to each branch (+*n*_*i*_ = +1,+2,+3 = 1bp, 2bp, 3bp insertions) and (−*n*_*i*_ = -1,-2,-3 = 1bp, 2bp, 3bp deletions), (8) we assigned a genotype to each phylogeny tip based on the sum of genotypes for each root-tip path to construct a vector of genotypes for tips in each phylogeny (**Figure S8**), (9) we computed Hs using TopDis (**Materials and Methods**) for each unique INDEL genotype in the vector for each phylogeny (i.e. *Hs*(−1) = 2, *Hs*(+1) = 2, *Hs*(+2) = 2 for the example phylogeny in **Figure S8**), (10) we aggregate Hs across all types of INDELs and for all eight phylogenies to get the Hs score for INDELs in the HT for a single simulation (*Hs*(*all INDELs in HT*) = 6 for phylogeny in **Figure S8**), (11) we repeat the steps above 1,000 times to get the probability that neutral INDELs arise in a 7bp HT ≥ *Hs* by taking the 99.8^th^ percentile from the distribution of *Hs* (all INDELs in HT) across all simulations which yields 45 INDELs within a HT and conclude that variation within HTs with ≥ 45 INDEL arisals in our dataset are unlikely to neutral.

### Homopolymeric Tract and Short Sequence Repeat Regions

We used the H37Rv reference genome to search for positions that corresponded to homopolymeric tracts and short sequence repeats in the Mtb genome. As phase variation has been documented with repeat units that consist between 1–7 nucleotides (Van Der Woude and Bäumler, 2004), we first classified regions with a single nucleotide repeated ≥ 7 times as homopolymeric tracts (HT) given the recent association in antibiotic tolerance (Safi et al., 2019; Vargas and Farhat, 2020) We scanned the genome for HTs at least 7bp in length and found 145 HT regions to cover 1,024bp or 0.023% of the genome (**Table S6**). Next, we searched the genome for regions in which a repeat unit, with any combination of nucleotides between 2-6bp, repeated at least 3 consecutive times (permutating four nucleotides for a 7bp unit yields too many possibilities to hold into memory). We classified these regions as short sequence repeats (SSR) and found them to cover 99,665bp or 2.26% of the genome.

### Association between Frameshifts in HTs and Antibiotic Resistance

In order to study the potential associations between the presence/absence of frameshift INDELs (relative to H37Rv) in specific HTs and antibiotic resistance, we used a publicly available dataset of antibiotic resistance phenotypic data (https://github.com/farhat-lab/resdata-ng/blob/master/resistance_data/summary_tables/resistance_summary.txt) (Gröschel et al., 2021). We determined the associations using a linear mixed model as implemented in GEMMA (Zhou and Stephens, 2012), allowing a maximum missingness of 1% (-miss parameter) and a minimum minor allele frequency of 1% (-maf parameter). In order to correct for population structure, we used a matrix of all SNP differences between the isolates tested. Finally, p-values were corrected for multiple testing using the Bonferroni method. For each test between frameshifts in a particular HT and antibiotic, we ensured we had ≥ 100 resistant isolates to that antibiotic in our sample.

### Gene Sets

Every gene on H37Rv was classified into one of six non-redundant gene categories according to the following criteria (Vargas et al., 2021): (i) genes identified as belonging to the PE/PPE family of genes unique to pathogenic mycobacteria, though to influence immunopathogenicity and characterized by conserved proline-glutamate (PE) and proline-proline-glutamate (PPE) motifs at the N protein termini (Brennan and Delogu, 2002; Comas et al., 2010; Phelan et al., 2016) were classified as *PE/PPE* (n = 167), (ii) genes flagged as being associated with antibiotic resistance (Farhat et al., 2013) were classified into the *Antibiotic Resistance* category (n = 28), (iii) genes encoding a CD4^+^ or CD8^+^ T-cell epitope (Comas et al., 2010; Coscolla et al., 2015) (but not already classified as a PE/PPE or Antibiotic Resistance gene) were classified as an *Antigen* (n = 257) , (iv) genes required for growth *in vitro* (Sassetti et al., 2003) and *in vivo* (Sassetti and Rubin, 2003) and not already placed into a category above were classified as *Essential* genes (n = 682), (v) genes flagged as transposases, integrases, phages or insertion sequences were classified as *Mobile Genetic Elements* (Comas et al., 2010) (n = 108), (vi) any remaining genes not already classified above were placed into the *Non-Essential* category (n = 2752).

### t-SNE Visualization

To construct the t-SNE plots that captured the genetic relatedness of the 31,428 isolates in our sample, we first constructed a pairwise SNP distance matrix. To efficiently compute this using our 782,565 x 31,428 genotypes matrix, we binarized the genotypes matrix and used sparse matrix multiplication implemented in Scipy to compute five 31,428 x 31,428 similarity matrices (Virtanen et al., 2020). We constructed a similarity matrix for each nucleotide (***A***, ***C***, ***G***, ***T***) where row *i*, column *j* of the similarity matrix for nucleotide *x* stored the number of *x*’s that isolate *i* and isolate *j* shared in common across all SNP sites. The fifth similarity matrix (***N***) stored the number of SNP sites in which neither isolate *i* and isolate *j* had a missing value. The pairwise SNP distance matrix (***D***) was then computed as ***D*** = ***N*** − (***A*** + ***C*** + ***G*** + ***T***). ***D*** had dimensions 31,428 x 31,428 where row *i*, column *j* stored the number of SNP sites in which isolate *i* and isolate *j* disagreed. We used ***D*** as input into a t-SNE algorithm implemented in Scikit-learn (Pedregosa et al., 2011) (settings: perplexity = 200, n_components = 2, metric = “precomputed”, n_iter = 1000, learning_rate = 2,500) to compute the embeddings for all 31,428 isolates in our sample. We used these embeddings to visualize the genetic relatedness of the isolates in two dimensions and colored isolates (points on the t-SNE plot) by lineage group (**Figure S4, Figure 3A**). For visualizing specific mutations, isolates were colored according to whether or not the alternate (mutant) allele was called (**Figure 3B-D**).

### Pathway Definitions

We used SEED (Overbeek et al., 2013) subsystem annotation to conduct pathway analysis and downloaded the subsystem classification for all features of *Mycobacterium tuberculosis* H37Rv (id: 83332.1) (Vargas et al., 2021). We mapped all of the annotated features from SEED to the annotation for H37Rv. Due to the slight inconsistency between the start and end chromosomal coordinates for features from SEED and our H37Rv annotation, we assigned a locus from H37Rv to a subsystem if both the start and end coordinates for this locus fell within a 20 base-pair window of the start and end coordinates for a feature in the SEED annotation (Vargas et al., 2021). We only included pathways that were composed of at least two genes.

### SNV and INDEL Mutational Density Calculation for Genes and Pathways

The homoplasy scores for all SNVs within each gene were aggregated to approximate all SNV mutation events (independent arisals) that occurred within the gene body then normalized by the gene length to calculate SNV mutational diversity for each gene (**Figure 5A**, **Table S8**). The homoplasy scores for all INDELs were computed similarly to approximate all INDEL mutation events then normalized by the gene length to calculate INDEL mutational diversity for each gene (**Figure 5B**, **Table S9**). When normalizing by gene length for both SNV and INDEL calculations, we removed positions with low Empirical Base Pair Recall scores (N=169,630), and excluded SNP sites: (**A**) missing calls in > 10% of isolates (N=31,215), (**B**) located in overlapping genes (N=933) (**Figure S2**). Further, we excluded genes that had an aggregate homoplasy score = 0 (no reported mutation events) and genes that were classified as Mobile Genetic Element for each set of computations (SNVs & INDELs). Next, we repeated our analysis at the level of pathways for SNVs (**Figure 5C**, **Table S10**) and INDELs (**Figure 5D**, **Table S11**) by aggregating mutations events occurring across genes belonging to each pathway and normalizing by the concatenate of the gene lengths. We again excluded positions with low Empirical Base Pair Recall scores (N=169,630), and excluded SNP sites: (**A**) missing calls in > 10% of isolates (N=31,215), (**B**) located in overlapping genes (N=933) when normalizing by the concatenate of gene lengths.

### Data Analysis and Variant Annotation

Data analysis was performed using custom scripts run in Python and interfaced with iPython (Pérez and Granger, 2007). Statistical tests were run with Statsmodels (Seabold and Perktold, 2010) and Figures were plotted using Matplotlib (Hunter, 2007). Numpy (Van Der Walt et al., 2011), Biopython (Cock et al., 2009) and Pandas (McKinney, 2010) were all used extensively in data cleaning and manipulation. Functional annotation of SNPs was done in Biopython using the H37Rv reference genome and the corresponding genome annotation. For every SNP variant called, we used the H37Rv reference position provided by the Pilon (Walker et al., 2014) generated VCF file to determine the nucleotide and codon positions if the SNP was located within a coding sequence in H37Rv. We extracted any overlapping CDS region and annotated SNPs accordingly, each overlapping CDS regions was then translated into its corresponding peptide sequence with both the reference and alternate allele. SNPs in which the peptide sequences did not differ between alleles were labeled synonymous, SNPs in which the peptide sequences did differ were labeled non-synonymous and if there were no overlapping CDS regions for that reference position, then the SNP was labeled intergenic. Functional annotation of indels was also done in Biopython using the H37Rv reference genome and the corresponding genome annotation. For every indel variant called, we used the H37Rv reference position provided by the Pilon generated VCF file to determine the nucleotide and codon positions if the indel was located within a coding sequence in H37Rv. An indel variant was classified as in-frame if the length of the indel allele was divisible by three, otherwise it was classified as a frameshift.

## DATA AND MATERIALS AVAILABILITY

Mtb sequencing data was collected from NCBI and is publicly available (**Materials and Methods**). All packages and software used in this study have been noted in the **Materials and Methods**. Custom scripts written in python version 2.7.15 were used to conduct all analyses and interfaced via Jupyter Notebooks. All scripts and notebooks will be uploaded to a GitHub repository upon acceptance of this manuscript for publication.

## AUTHOR CONTRIBUTIONS

R.V.J., and M.R.F. conceived the idea for the study. M.L., C.S., and K.M. conceived the idea for the fluctuation analysis and contributed to the fluctuation analysis. M.R.F., and C.S. supervised the project. R.V.J. performed data acquisition, data curation, and data analysis. M.L. carried out fluctuation analysis experiments. L.F. curated data and performed the GWAS analysis. M.L., and T.I. carried out differential expression experiments and analysis for *espA* mutants. R.V.J. and M.R.F. wrote the first draft. M.R.F. and C.S. critically reviewed the drafts. All authors reviewed the draft and assisted in the manuscript preparation.

## ACKNOWLEDGEMENTS

We thank the members of the Farhat lab for helpful discussions and comments on the research project and manuscript. R.V.J. was supported by the National Science Foundation Graduate Research Fellowship under Grant No. DGE1745303. M.F. was supported by NIH NIAID R01 AI55765. Portions of this research were conducted on the O2 High Performance Compute Cluster, supported by the Research Computing Group, at Harvard Medical School.

## SUPPLEMENTARY DATA SUPPLEMENTARY FIGURES

**Figure S1.**
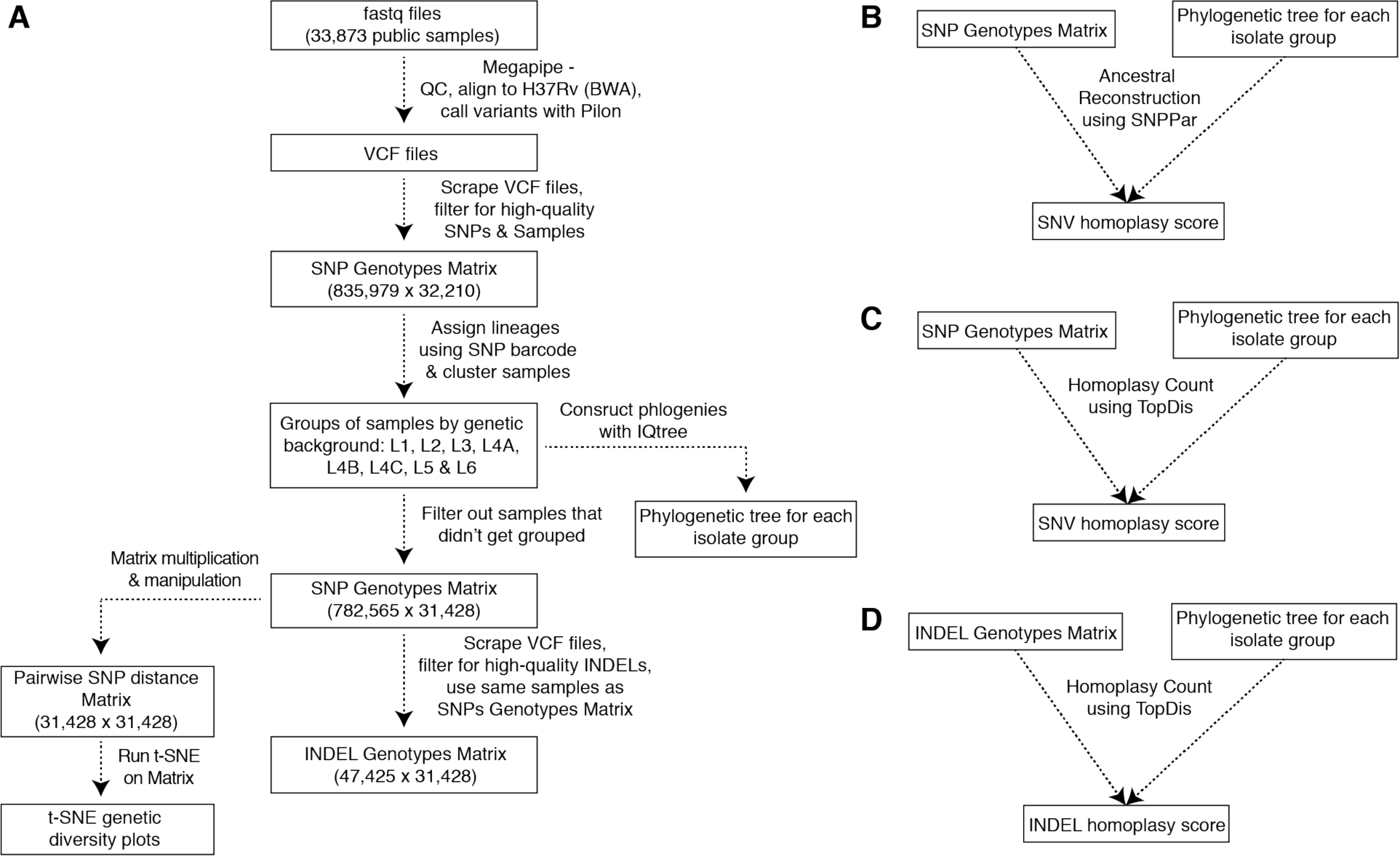
Project Workflow. (**A**) We processed 33,873 public sequences. After quality-control steps removed low-quality SNP sites and isolates, and we removed isolates that did not get classified into one of eight lineage groups (L1, L2, L3, L4A, L4B, L4C, L5, L6) we constructed a 782,565x31,428 SNP genotypes matrix (**Figure S2**). We used this SNP genotypes matrix to construct a pairwise SNP distance matrix which was then used as an input into a t-SNE algorithm. We scraped the VCF files for 31,428 isolates in the SNP genotypes matrix to construct a 47,425x31,428 INDEL genotypes matrix. We constructed a phylogeny for each lineage group (**Materials and Methods**). (**B**) Homoplasy Scores for SNVs were computed using SNPPar with the SNP genotypes matrix and phylogenies as input. (**C**) Homoplasy Scores for SNVs were computed using TopDis with the SNP genotypes matrix and phylogenies as input. (**D**) Homoplasy Scores for INDELs were computed using TopDis with the INDEL genotypes matrix and phylogenies as input.

**Figure S2.**
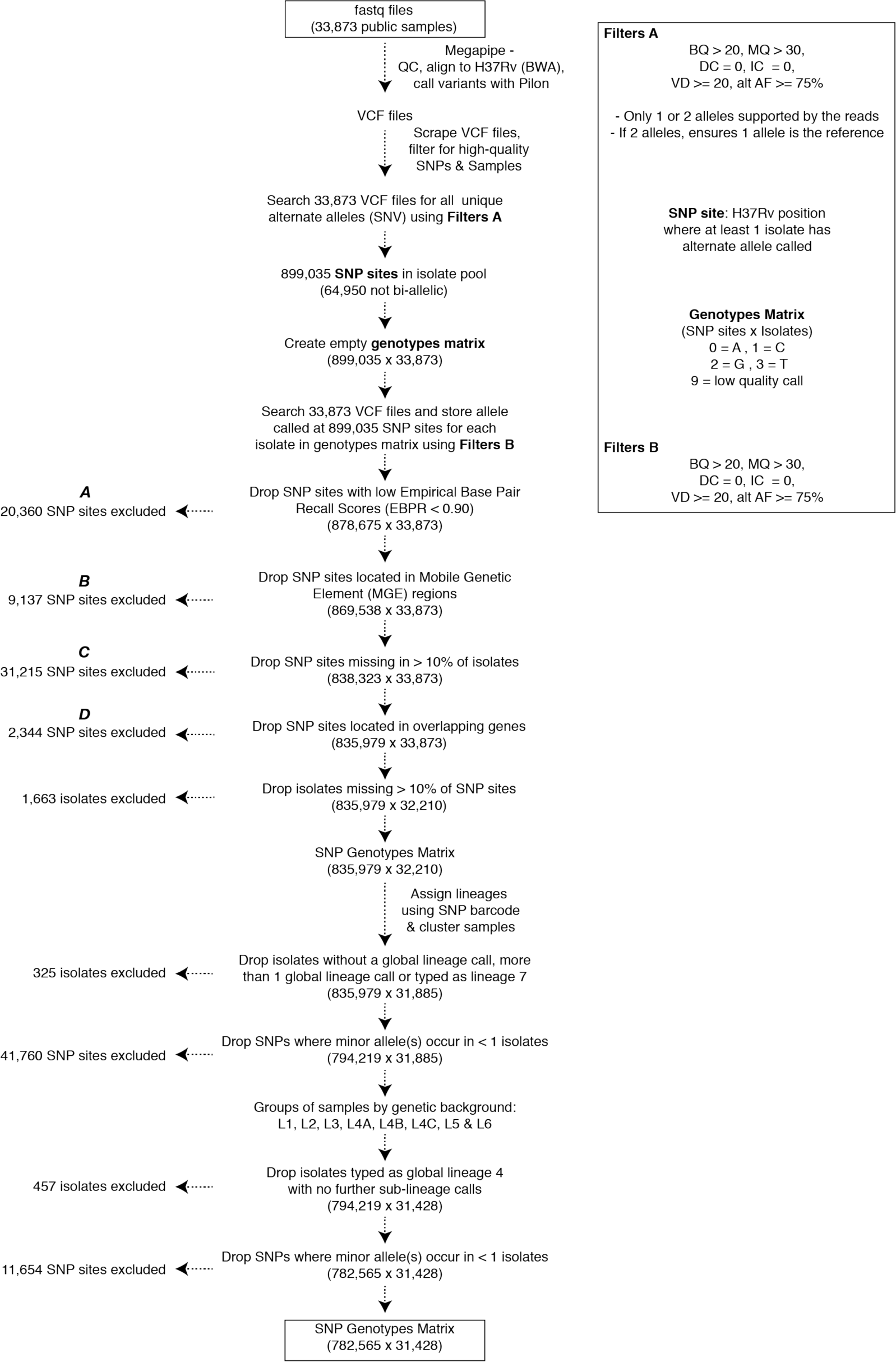
Constructing the SNP genotypes matrix. A schematic diagram outlining the steps described in **Materials and Methods/SNP Genotypes Matrix** and relevant QC filters; from downloading public sequences to creating the final SNP genotypes matrix.

**Figure S3.**
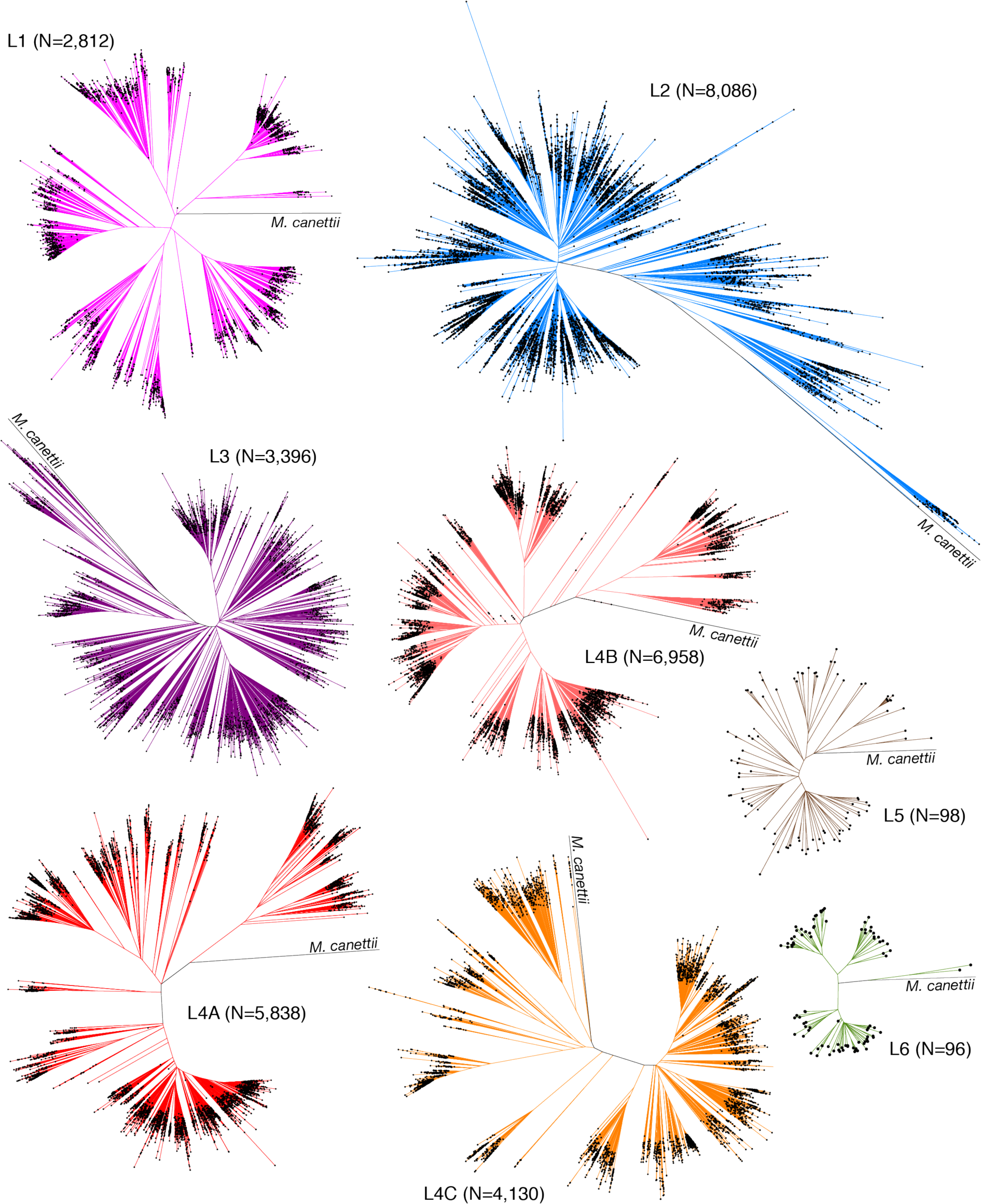
Maximum likelihood phylogenies for 31,428 isolates. We separated 31,428 isolates into eight groups by genetic background and constructed eight separate phylogenies. Misclassified isolates were pruned from the phylogenies for visualization (14 isolates total: L1(3), L2(4), L3(2), L4A(1), L4C(4)) (N = # of isolates).

**Figure S4.**
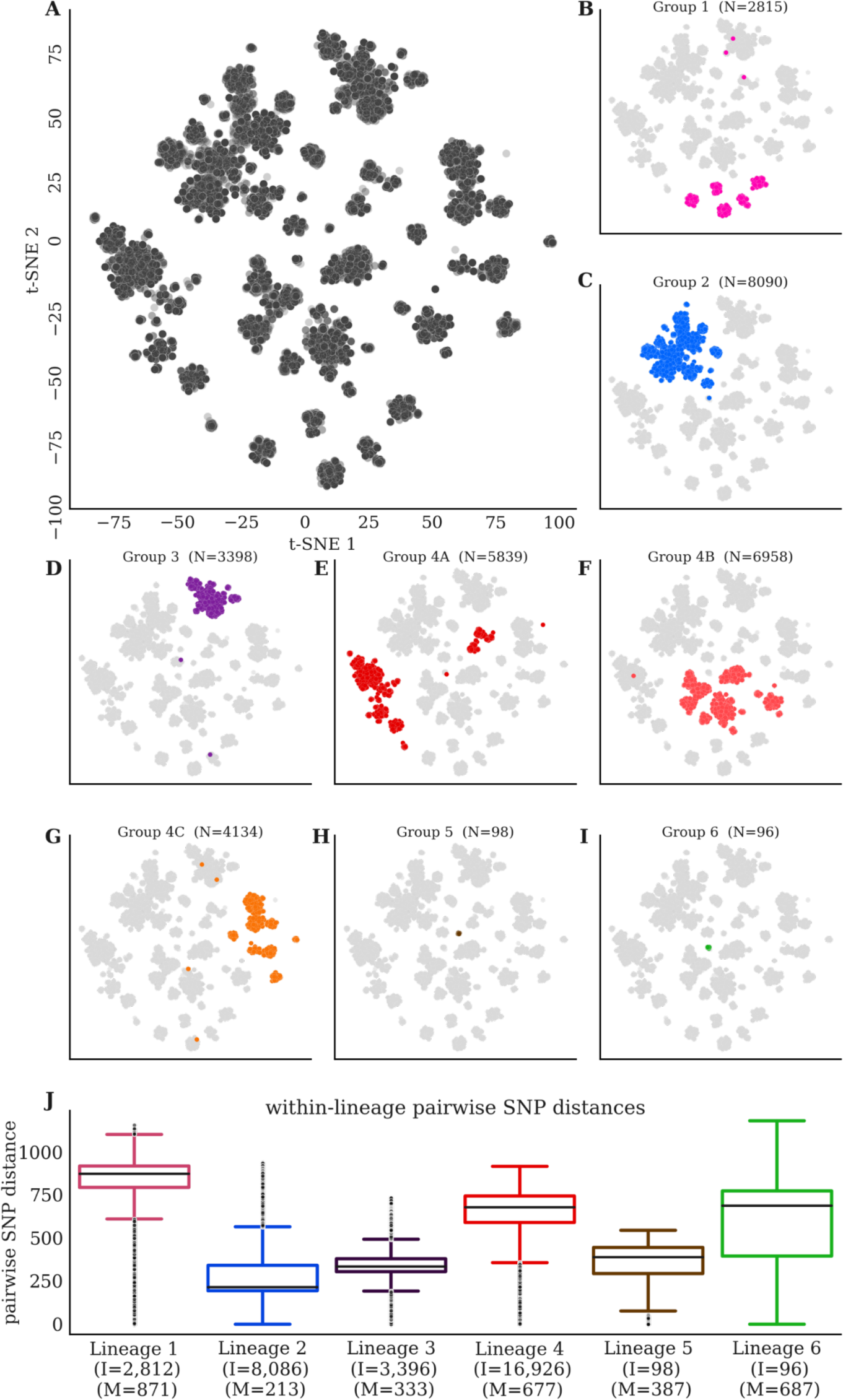
Genetic distance between 31,428 samples separates lineages. (**A**) t-SNE plots of pairwise SNP distance across the global sample of 31,428 clinical isolates and 782,565 SNP sites. (**B-I**) Isolates in the t-SNE colored by lineage/sub-lineage (L1, L2, L3, L4A, L4B, L4C, L5, L6) (N = # of isolates). (**J**) Pairwise SNP distances between each pair of isolates within each lineage L1-L6. The 14/31,328 misclassified isolates were removed prior to computing the distribution of pairwise distances for these barplots (N = # of isolates, M = median of pairwise SNP distances).

**Figure S5.**
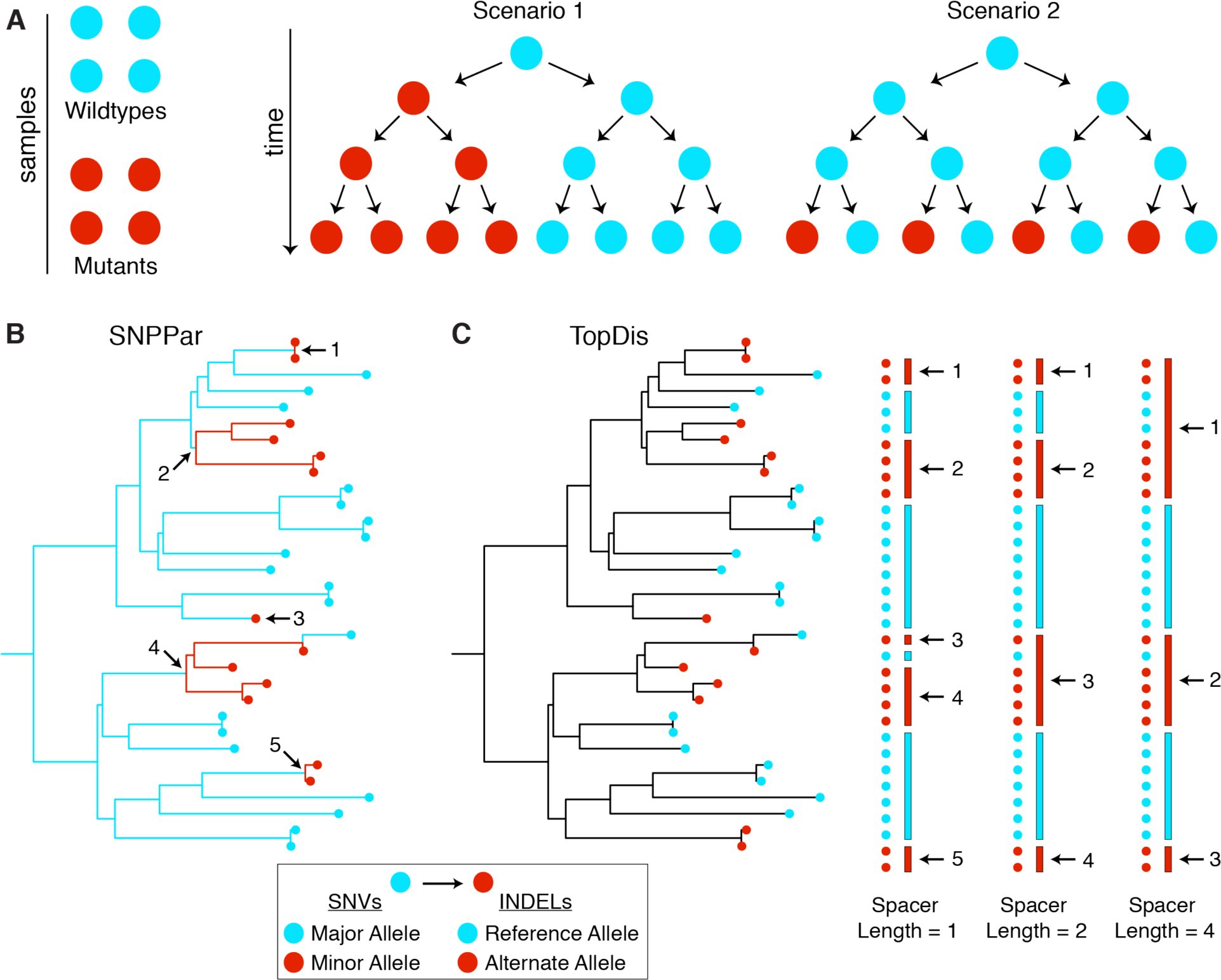
SNPPar and TopDis methods for computing homoplasy scores. Phylogenies are required to discern how many times a given mutation arose within a pool of samples. (**A**) In scenario 1, the mutation arose once in an ancestor while in scenario 2 the mutation arose independently on four occasions which providing much stronger evidence that this mutation was a target of positive selection. (**B**) SNPPar is the ancestral reconstruction program we used to infer where SNV mutations occurred on the trees and consequently how many times a mutation *arose* in the tree. (**C**) TopDis is our own method of counting the number of *mutation events* for a given variant. TopDis includes a parameter (Spacer Length) that controls how conservatively we count a single independent mutation given the topology of the tree (**Materials and Methods**).

**Figure S6.**
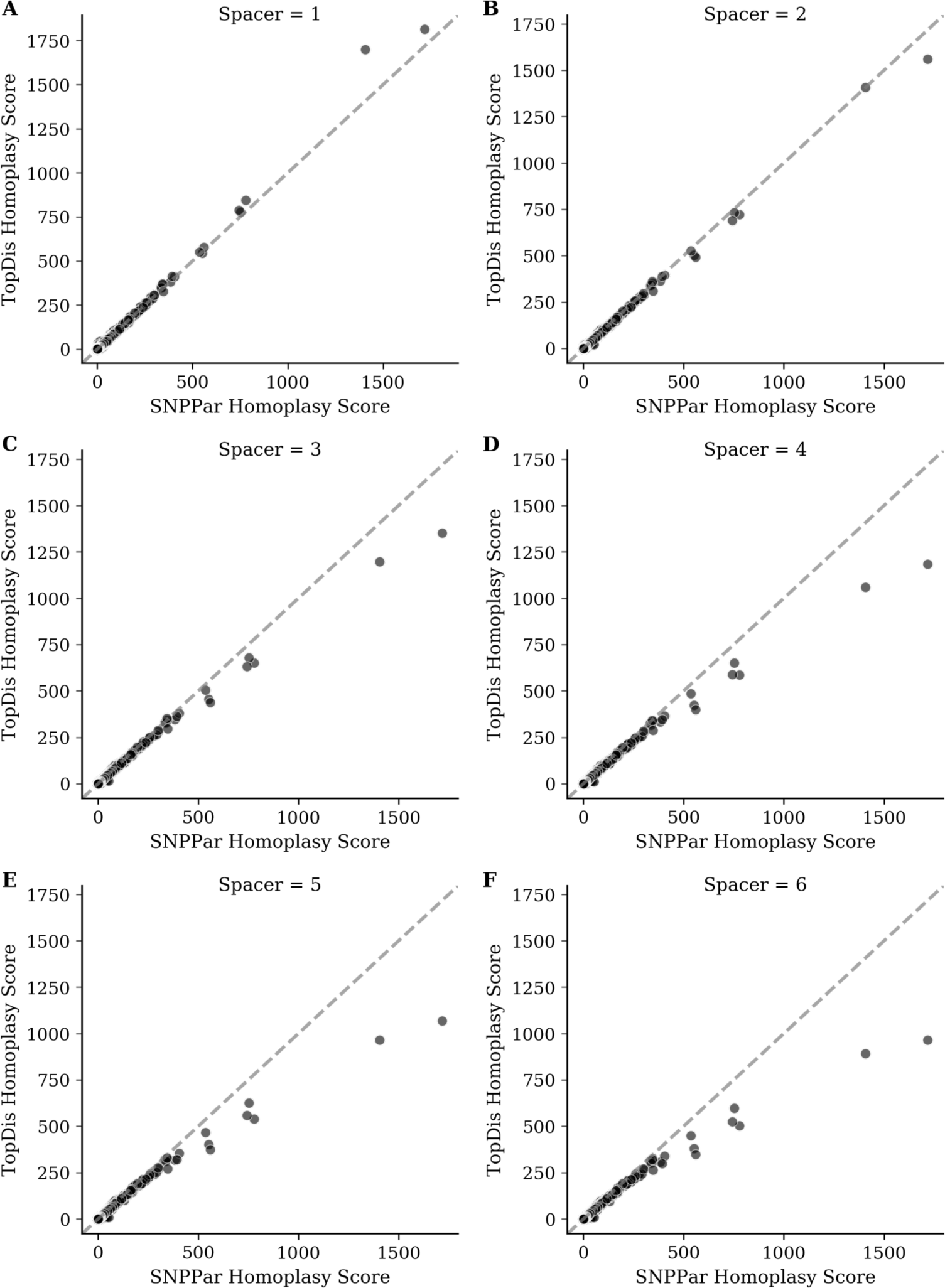
Homoplasy scores from SNPPar vs. TopDis. We calculated homoplasy scores for all 836,901 SNVs in our dataset using SNPPar (**Figure S5B**) and TopDis (**Figure S5C**). TopDis takes in a parameter (Spacer) which calculates more conservative estimates for the number of independent arisals for larger values (**Figure S5C**, **Materials and Methods**). (**A**-**F**) We compare homoplasy scores obtained from SNPPar to those obtained from TopDis for different values of Spacer (1, 2, 3, 4, 5, 6). The two methods obtain similar estimates for the number of independent arisals across the phylogenies, with TopDis yielding lower esimates than SNPPar for larger values of the Spacer parameter. We conservatively chose TopDis Spacer = 4 for INDEL homoplasy score calculations.

**Figure S7.**
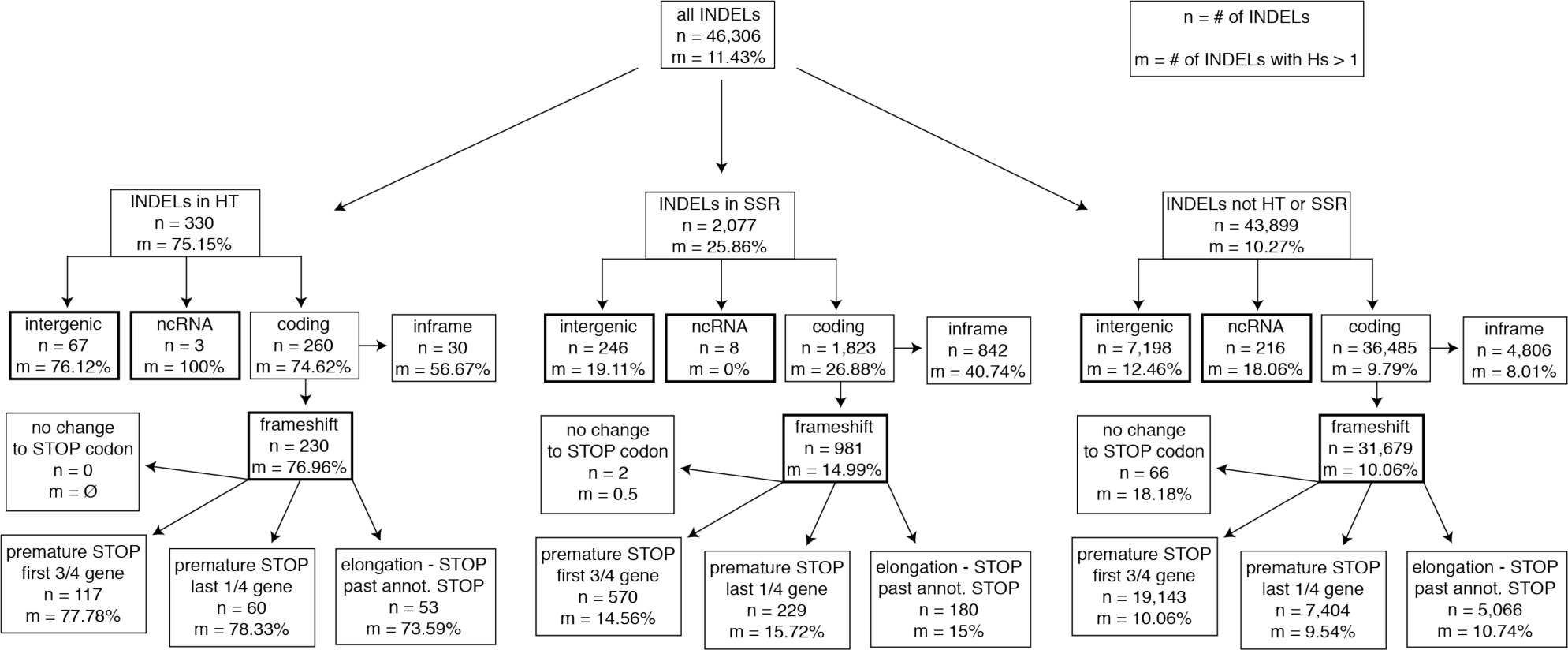
Functional breakdown of INDELs for HT, SSR & other regions of the genome. We detected 46,306 INDEL genotypes in our sample, of these 11.43% were homoplastic (Hs > 1) and independently arose more than once in our sample. Breaking down INDELs by whether they occur in HT regions, SSR regions or other regions of the genome reveals a substantially higher proportion of INDELs that are homoplastic in HT regions (75.15%) than in SSR regions (25.86%) and other regions (10.27%). Further breakdown of INDELs by functional impact shows a high proportion of homoplastic variants among frameshifts detected within HT regions (76.96%) relative to frameshifts in SSR regions (14.99%) and frameshifts in other regions (10.06%).

**Figure S8.**
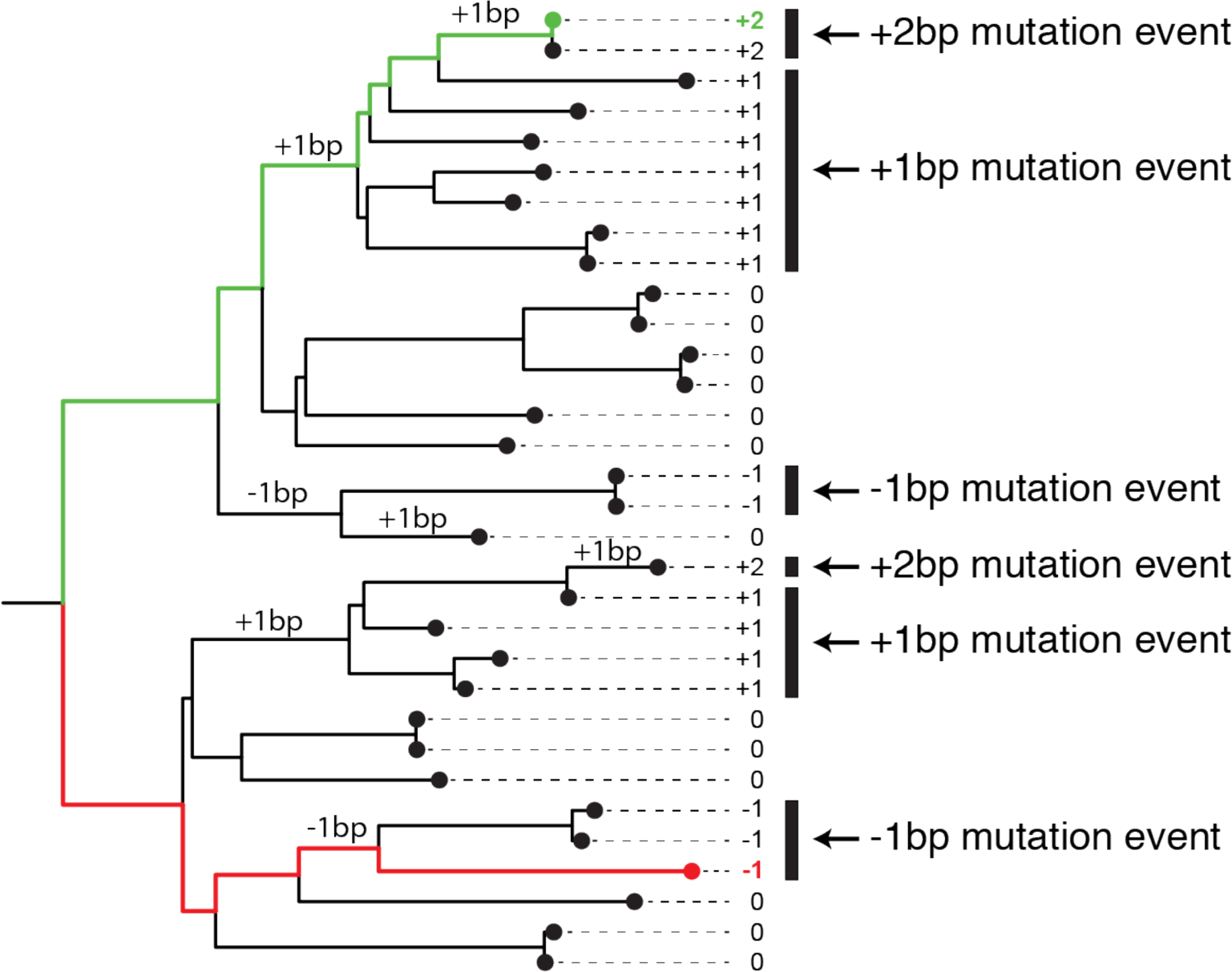
Simulations for frameshifts in HT regions under neutrality. A schematic demonstrating how simulations of neutral frameshift mutations in a HT were carried out using the phylogenies generated from our sample (**Methods**). Briefly, (1) 1-bp frameshift mutations were simulated on the branches of a given phylogeny, (2) INDEL genotypes were assigned for each tip (the green path results in a 2-bp insertion resulting from two 1-bp frameshift insertions in the HT occurring along the path from the root to the tip, the red path results in a 1-bp deletion at the tip), (3) TopDis is used to calculate Hs for each unique INDEL genotype represented at the tips: Hs(+2bp) = 2, Hs(+1bp) = 2, Hs(-1bp) = 2, (4) Hs is aggregated for all genotypes across all phylogenies to get a cumulative Hs for a HT, Hs(HT) = 6 for example above. This process is repeated 1000 times to create a null distribution of Hs (HT) under neutrality for the phylogenies used in our analyses.

**Figure S9.**
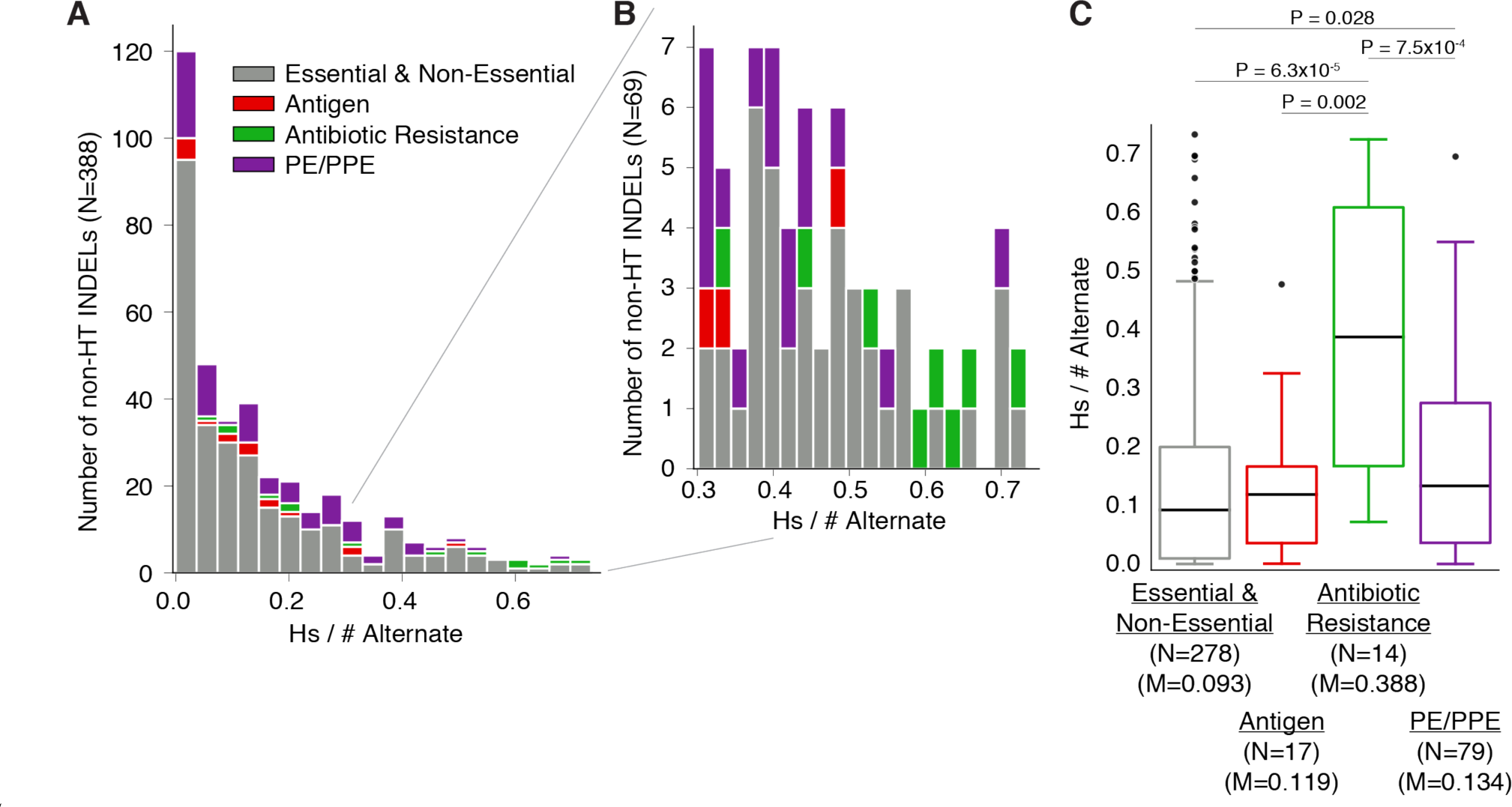
Recency Ratio for non-HT INDELs. (**A-B**) The distribution of the ratio of (homoplasy score) to (# of isolates harboring the alternate allele) for 388/655 INDELs (**Figure 1C**) that occur in non-HT (SSR & other) and coding regions. (**C**) Breaking these INDEL ratios down by gene category reveals higher ratios overall for antibiotic resistance genes when compared to other gene categories. N = number of alleles, M = median RcR

## SUPPLEMENTARY TABLES

**Table S1.** **SNVs with the 30 highest homoplasy scores.** The *# Minor* column contains the number of isolates harboring the minor allele in our sample of 31,428 isolates. The recency ratio (RcR = *Homoplasy Score* / *# Minor*) is given in the rightmost column. Table S2 lists all 1,525 SNVs with homoplasy score ≥ 5 and minor allele frequency > 0.1% and the breakdown of mutation arisals by lineage.

| H37Rv<br>Position | Ref | Alt | Minor | Gene Symbol | Gene<br>Coord | AA<br>change | Homoplasy<br>Score | # Minor | RcR |
| --- | --- | --- | --- | --- | --- | --- | --- | --- | --- |
| 2155168 | C | G | G | *katG* | 944 | S315T | 1717 | 8066 | 0.21 |
| 761155 | C | T | T | *rpoB* | 1349 | S450L | 1406 | 5706 | 0.25 |
| 781687 | A | G | G | *rpsL* | 128 | K43R | 779 | 3509 | 0.22 |
| 1673425 | C | T | T | *Rv1482c-fabG1* |  |  | 752 | 2660 | 0.28 |
| 4247429 | A | G | G | *embB* | 916 | M306V | 743 | 2102 | 0.35 |
| 1473246 | A | G | G | *Rrs* | 1401 |  | 560 | 1452 | 0.39 |
| 7582 | A | G | G | *gyrA* | 281 | D94G | 552 | 844 | 0.65 |
| 4247431 | G | A | A | *embB* | 918 | M306I | 536 | 1208 | 0.44 |
| 3883626 | A | G | G | *Rv3466* | 102 | P34P | 405 | 9033 | 0.05 |
| 3884906 | A | G | A | *Rv3467* | 943 | E315K | 393 | 6211 | 0.06 |
| 7570 | C | T | T | *gyrA* | 269 | A90V | 384 | 596 | 0.64 |
| 3730411 | G | A | A | *PPE54* | 6525 | G2175<br>G | 347 | 7494 | 0.05 |
| 105060 | G | A | A | *Rv0095c* | 156 | D52D | 345 | 2066 | 0.17 |
| 1164571 | A | G | G | *PE8-Rv1041c* |  |  | 343 | 7811 | 0.04 |
| 608037 | A | C | C | *Rv0515* | 1487 | H496P | 337 | 5235 | 0.06 |
| 105063 | G | A | A | *Rv0095c* | 153 | F51F | 333 | 2080 | 0.16 |
| 401678 | C | A | A | *Rv0336* | 1487 | P496H | 301 | 1796 | 0.17 |
| 3136335 | G | A | A | *Rv2828c-*<br>*vapC22* |  |  | 299 | 3907 | 0.08 |
| 2123145 | C | T | T | *lldD2* | 7 | V3I | 291 | 1953 | 0.15 |
| 761110 | A | T | T | *rpoB* | 1304 | D435V | 279 | 802 | 0.35 |
| 781822 | A | G | G | *rpsL* | 263 | K88R | 276 | 843 | 0.33 |
| 761139 | C | T | T | *rpoB* | 1333 | H445Y | 260 | 429 | 0.61 |
| 2626600 | G | A | A | *esxP-Rv2348c* |  |  | 258 | 5158 | 0.05 |
| 1094538 | T | G | G | *PE_PGRS17-*<br>*Rv0979c* |  |  | 254 | 1357 | 0.19 |
| 2122395 | C | T | T | *lldD2* | 757 | V253M | 254 | 10090 | 0.03 |
| 1096633 | T | G | T | *PE_PGRS18-*<br>*mprA* |  |  | 241 | 1212 | 0.2 |
| 1276588 | C | G | G | *Rv1148c* | 1161 | A387A | 239 | 12568 | 0.02 |
| 4248003 | A | G | G | *embB* | 1490 | Q497R | 239 | 490 | 0.49 |
| 1339399 | C | T | T | *PPE18* | 51 | Y17Y | 226 | 2416 | 0.09 |
| 2439204 | A | G | G | *pknL-Rv2177c* |  |  | 223 | 1004 | 0.22 |

Table S2**. Homoplasy scores for 1,525 SNVs.** A full version of Table S1. Homoplasy scores for 1,525 SNVs with homoplasy score ≥ 5 and minor allele frequency > 0.1% computed from the ancestral reconstruction method (SNPPar). The *Homoplasy Score* column contains the number of inferred independent arisals aggregated across all of the phylogenies. The *# Minor* column contains the number of isolates harboring the minor allele in our sample of 31,428 isolates. The *Homoplasy Score* / *# Minor* contains the ratio of these two columns. Columns *L1*, *L2*, *L3*, *L4A*, *L4B*, *L4C*, *L5*, *L6* correspond to number independent arisals broken down by phylogeny. (Excel spreadsheet)

**Table S3.**
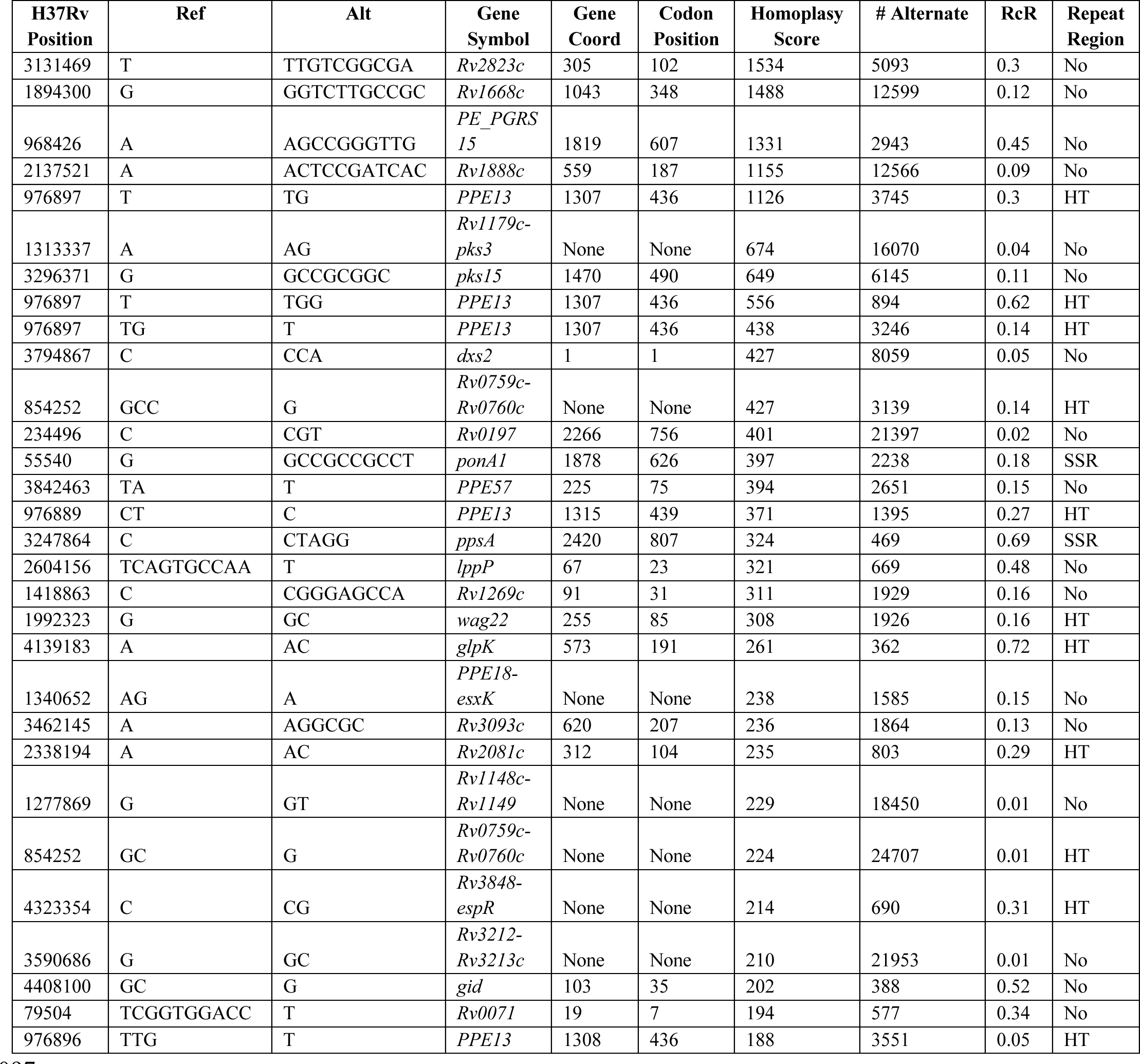
**INDELs with the 30 highest homoplasy scores.** The *# Alternate* column contains the number of isolates harboring the alternate allele in our sample of 31,428 isolates. The recency ratio (RcR = *Homoplasy Score* / *# Alternate*) is given in the rightmost column. Table S4 lists all 655 INDELs with homoplasy score ≥ 5 and alternate allele frequency > 0.1% and the breakdown of mutation arisals by lineage.

Table S4**. Homoplasy scores for 655 INDELs.** A full version of Table S3. Homoplasy scores for 655 INDELs with homoplasy score ≥ 5 and alternate allele frequency > 0.1% computed from the naïve phylogeny counting method (TopDis). The *Homoplasy Score* column contains the number of inferred independent arisals aggregated across all of the phylogenies. The *# Alternate* column contains the number of isolates harboring the alternate allele in our sample of 31,428 isolates. The *Homoplasy Score* / *# Alternate* contains the ratio of these two columns. Columns *L1*, *L2*, *L3*, *L4A*, *L4B*, *L4C*, *L5*, *L6* correspond to number independent arisals broken down by phylogeny. (Excel spreadsheet)

**Table S5.** **Homopolymeric tracts with more than 45 independent frameshift arisals.** The frameshift variants occurring within these HT regions have an aggregated homoplasy score of > 45. The *Homoplasy Score* column lists the number of independent frameshift mutation arisals in the corresponding HT for HTs in coding regions and number of independent INDEL arisals for HTs in intergenic regions. The *# Alternate* column gives the number of isolates that harbor a frameshift alternate allele in this HT. These HT regions are highlighted in blue in **Figure 1C** and some mutations within the HTs for the bolded rows are represented in **Figure 3B-D**. A full version of this table that lists all 145 HT regions found in MTBC genome can be found in Table S6.

| H37Rv Start | H37Rv End | polyNT | Gene Symbol | sum(Hs) | # Alternate |
| --- | --- | --- | --- | --- | --- |
| 976897 | 976906 | GGGGGGGGG | *PPE13* | 2317 | 8351 |
| 854252 | 854261 | CCCCCCCCC | *Rv0759c-Rv0760c* | 776 | 28077 |
| 976889 | 976896 | TTTTTTTTT | *PPE13* | 771 | 5641 |
| 1992323 | 1992331 | CCCCCCCCC | *wag22* | 578 | 4052 |
| 2338194 | 2338202 | CCCCCCCCC | *Rv2081c* | 360 | 4596 |
| **4139183** | **4139190** | **CCCCCCCC** | ***glpK*** | **282** | **388** |
| 1333661 | 1333668 | GGGGGGGG | *Rv1190* | 277 | 914 |
| 4323354 | 4323361 | GGGGGGGG | *Rv3848-espR* | 269 | 1355 |
| 2234247 | 2234254 | GGGGGGGG | *Rv1990c-mazF6* | 217 | 901 |
| 364498 | 364505 | GGGGGGGG | *vapC2-Rv0302* | 216 | 676 |
| 3742991 | 3742998 | GGGGGGGG | *PE_PGRS50-Rv3346c* | 214 | 2229 |
| 691887 | 691894 | CCCCCCCC | *mce2D* | 198 | 5158 |
| **4358979** | **4358986** | **GGGGGGGG** | ***espK*** | **192** | **902** |
| 1760164 | 1760171 | GGGGGGGG | *frdB* | 178 | 791 |
| 4099402 | 4099409 | GGGGGGGG | *B11* | 163 | 656 |
| 1700415 | 1700422 | GGGGGGGG | *Rv1509* | 153 | 399 |
| 2976558 | 2976565 | CCCCCCCC | *Rv2652c-Rv2653c* | 152 | 669 |
| 36470 | 36477 | CCCCCCCC | *bioF2* | 140 | 4903 |
| 2536625 | 2536632 | GGGGGGGG | *Rv2264c* | 138 | 8974 |
| 3308313 | 3308320 | GGGGGGGG | *Rv2955c* | 137 | 472 |
| 4337820 | 4337827 | GGGGGGGG | *Rv3860* | 131 | 337 |
| **4056480** | **4056487** | **TTTTTTTT** | ***espA-ephA*** | **108** | **405** |
| 2796131 | 2796138 | CCCCCCCC | *PE_PGRS42* | 93 | 762 |
| 1026916 | 1026923 | GGGGGGGG | *Rv0920c-Rv0921* | 90 | 332 |
| 2881582 | 2881589 | TTTTTTTT | *Rv2560-Rv2563* | 89 | 3701 |
| 799136 | 799143 | CCCCCCCC | *Rv0698* | 83 | 4229 |
| 2522481 | 2522488 | GGGGGGGG | *Rv2248* | 79 | 250 |
| 191391 | 191398 | CCCCCCCC | *Rv0161* | 73 | 274 |
| 2141408 | 2141415 | GGGGGGGG | *Rv1894c* | 72 | 433 |
| 1002282 | 1002289 | CCCCCCCC | *Rv0897c* | 69 | 294 |
| 2440187 | 2440194 | GGGGGGGG | *Rv2177c-aroG* | 69 | 819 |
| 1572680 | 1572687 | CCCCCCCC | *PE_PGRS25* | 68 | 1792 |
| 4325205 | 4325212 | AAAAAAA | *Hns* | 64 | 155 |
| 868160 | 868167 | GGGGGGGG | *Rv0774c* | 64 | 151 |
| 3112277 | 3112284 | CCCCCCCC | *Rv2803* | 63 | 130 |
| 4019546 | 4019553 | GGGGGGGG | *Rv3577* | 63 | 149 |
| 3539139 | 3539146 | GGGGGGGG | *aofH* | 62 | 2862 |
| 3602343 | 3602350 | CCCCCCCC | *Rv3225c* | 62 | 157 |
| 4251293 | 4251300 | GGGGGGGG | *fadE35* | 59 | 142 |
| 1546465 | 1546472 | CCCCCCCC | *Rv1373* | 58 | 333 |
| 2001789 | 2001796 | GGGGGGGG | *PE_PGRS31* | 57 | 361 |
| 3559990 | 3559997 | CCCCCCCC | *Rv3192* | 55 | 152 |
| 3377346 | 3377353 | CCCCCCCC | *PPE46* | 54 | 116 |
| 2867737 | 2867744 | CCCCCCCC | *lppB* | 50 | 262 |
| 4298220 | 4298227 | GGGGGGGG | *pks2* | 48 | 96 |

**Table S6. Homoplasy scores aggregated by homopolymeric tracts.** A full version of Table S5. This table lists all 145 HT regions found in the MTBC genome. For each HT, the homoplasy scores for all frameshifts occurring within HTs in coding regions and homoplasy scores for all INDELs occurring in intergenic regions are aggregated which yields a *Homoplasy Score* for each HT. The *# Alternate* column gives the number of isolates that harbor a frameshift alternate allele in this HT. (Excel spreadsheet)

**Table S7. DeSeq analysis.** Results from DeSeq analyses showing 22 significantly differentially expressed genes in *espA* mutant. (Excel spreadsheet)

**Table S8. SNV mutational density per gene.** The homoplasy scores for all SNVs within each gene were aggregated to approximate all SNV mutation events (independent arisals) that occurred within the gene body then normalized by the gene length. This table contains these calculations for each gene as well as columns for *# SNVs*, *Synonymous Homoplasy Score*, and *Non-Synonymous Homoplasy Score*. (Excel spreadsheet)

**Table S9. INDEL mutation density per gene.** The homoplasy scores for all INDELs within each gene were aggregated to approximate all INDEL mutation events (independent arisals) that occurred within the gene body then normalized by the gene length. This table contains these calculations for each gene as well as columns for *# INDELs*, *Inframe Homoplasy Score*, and *Frameshift Homoplasy Score*. (Excel spreadsheet)

**Table S10. SNV mutational density per pathway.** The homoplasy scores for all SNVs within each pathway were aggregated to approximate all SNV mutation events (independent arisals) that occurred within the genes in each pathway then normalized by the concatenate of the gene lengths. This table contains these calculations for each pathway as well as columns for *H37Rv Locus Tags* and *Gene Symbols* for the genes that belong to each pathway. (Excel spreadsheet)

**Table S11. INDEL mutation density per pathway.** The homoplasy scores for all INDELs within each pathway were aggregated to approximate all INDEL mutation events (independent arisals) that occurred within the genes in each pathway then normalized by the concatenate of the gene lengths. This table contains these calculations for each pathway as well as columns for *H37Rv Locus Tags* and *Gene Symbols* for the genes that belong to each pathway. (Excel spreadsheet)

## REFERENCES

1. Bellerose MM, Baek S-H, Huang C-C, Moss CE, Koh E-I, Proulx MK, Smith CM, Baker RE, Lee JS, Eum S. 2019. Common Variants in the Glycerol Kinase Gene Reduce Tuberculosis Drug Efficacy. mBio 10:e00663–19.

2. Benson DA, Karsch-Mizrachi I, Lipman DJ, Ostell J, Sayers EW. 2000. GenBank. Nucleic Acids Res 28:15–18.

3. Brennan MJ. 2017. The enigmatic PE/PPE multigene family of mycobacteria and tuberculosis vaccination. Infect Immun 85:e00969–16.

4. Brennan MJ, Delogu G. 2002. The PE multigene family: a ‘molecular mantra’for mycobacteria. Trends Microbiol 10:246–249.

5. Brynildsrud OB, Pepperell CS, Suffys P, Grandjean L, Monteserin J, Debech N, Bohlin J, Alfsnes K, Pettersson JO-H, Kirkeleite I. 2018. Global expansion of Mycobacterium tuberculosis lineage 4 shaped by colonial migration and local adaptation. Sci Adv 4:eaat5869.

6. Chiner-Oms Á, Sánchez-Busó L, Corander J, Gagneux S, Harris S, Young D, González-Candelas F, Comas I. 2019. Genomic determinants of speciation and spread of the Mycobacterium tuberculosis complex. Sci Adv 5:eaaw3307.

7. Clemmensen HS, Knudsen NPH, Rasmussen EM, Winkler J, Rosenkrands I, Ahmad A, Lillebaek T, Sherman DR, Andersen PL, Aagaard C. 2017. An attenuated Mycobacterium tuberculosis clinical strain with a defect in ESX-1 secretion induces minimal host immune responses and pathology. Sci Rep 7:1–13.

8. Cock PJA, Antao T, Chang JT, Chapman BA, Cox CJ, Dalke A, Friedberg I, Hamelryck T, Kauff F, Wilczynski B, others. 2009. Biopython: freely available Python tools for computational molecular biology and bioinformatics. Bioinformatics 25:1422–1423.

9. Cole ST. 2016. Inhibiting Mycobacterium tuberculosis within and without. Philos Trans R Soc B Biol Sci 371:20150506.

10. Coll F, Phelan J, Hill-Cawthorne GA, Nair MB, Mallard K, Ali S, Abdallah AM, Alghamdi S, Alsomali M, Ahmed AO. 2018. Genome-wide analysis of multi-and extensively drug-resistant Mycobacterium tuberculosis. Nat Genet 50:307.

11. Comas I, Chakravartti J, Small PM, Galagan J, Niemann S, Kremer K, Ernst JD, Gagneux S. 2010. Human T cell epitopes of Mycobacterium tuberculosis are evolutionarily hyperconserved. Nat Genet 42:498–503.

12. Coscolla M, Copin R, Sutherland J, Gehre F, de Jong B, Owolabi O, Mbayo G, Giardina F, Ernst JD, Gagneux S. 2015. M. tuberculosis T cell epitope analysis reveals paucity of antigenic variation and identifies rare variable TB antigens. Cell Host Microbe 18:538–548.

13. Coscolla M, Gagneux S. 2014. Consequences of genomic diversity in Mycobacterium tuberculosis. Presented at the Seminars in immunology. Elsevier. pp. 431–444.

14. Coscolla M, Gagneux S, Menardo F, Loiseau C, Ruiz-Rodriguez P, Borrell S, Otchere ID, Asante-Poku A, Asare P, Sánchez-Busó L. 2021. Phylogenomics of Mycobacterium africanum reveals a new lineage and a complex evolutionary history. Microb Genomics 7:000477.

15. Covert BA, Spencer JS, Orme IM, Belisle JT. 2001. The application of proteomics in defining the T cell antigens of Mycobacterium tuberculosis. PROTEOMICS Int Ed 1:574–586.

16. DeJesus MA, Gerrick ER, Xu W, Park SW, Long JE, Boutte CC, Rubin EJ, Schnappinger D, Ehrt S, Fortune SM. 2017. Comprehensive essentiality analysis of the Mycobacterium tuberculosis genome via saturating transposon mutagenesis. MBio 8:e02133–16.

17. Edwards DJ, Duchêne S, Pope B, Holt KE. 2020. SNPPar: identifying convergent evolution and other homoplasies from microbial whole-genome alignments. *bioRxiv*.

18. Ektefaie Y, Dixit A, Freschi L, Farhat MR. 2021. Globally diverse Mycobacterium tuberculosis resistance acquisition: a retrospective geographical and temporal analysis of whole genome sequences. Lancet Microbe 2:e96–e104.

19. Estrem ST, Ross W, Gaal T, Chen ZS, Niu W, Ebright RH, Gourse RL. 1999. Bacterial promoter architecture: subsite structure of UP elements and interactions with the carboxy-terminal domain of the RNA polymerase α subunit. Genes Dev 13:2134– 2147.

20. Farhat MR, Shapiro BJ, Kieser KJ, Sultana R, Jacobson KR, Victor TC, Warren RM, Streicher EM, Calver A, Sloutsky A, others. 2013. Genomic analysis identifies targets of convergent positive selection in drug-resistant Mycobacterium tuberculosis. Nat Genet 45:1183–1183.

21. Fortune S, Jaeger A, Sarracino D, Chase M, Sassetti C, Sherman D, Bloom B, Rubin E. 2005. Mutually dependent secretion of proteins required for mycobacterial virulence. Proc Natl Acad Sci 102:10676–10681.

22. Freschi L, Vargas R, Hussain A, Kamal SM, Skrahina A, Tahseen S, Ismail N, Barbova A, Niemann S, Cirillo DM. 2021. Population structure, biogeography and transmissibility of Mycobacterium tuberculosis. Nat Commun **In-press**.

23. Gagneux S. 2018. Ecology and evolution of Mycobacterium tuberculosis. Nat Rev Microbiol 16:202–213.

24. Garces A, Atmakuri K, Chase MR, Woodworth JS, Krastins B, Rothchild AC, Ramsdell TL, Lopez MF, Behar SM, Sarracino DA. 2010. EspA acts as a critical mediator of ESX1- dependent virulence in Mycobacterium tuberculosis by affecting bacterial cell wall integrity. PLoS Pathog 6:e1000957.

25. Gerrick ER, Barbier T, Chase MR, Xu R, François J, Lin VH, Szucs MJ, Rock JM, Ahmad R, Tjaden B. 2018. Small RNA profiling in Mycobacterium tuberculosis identifies MrsI as necessary for an anticipatory iron sparing response. Proc Natl Acad Sci 115:6464– 6469.

26. Gill WP, Harik NS, Whiddon MR, Liao RP, Mittler JE, Sherman DR. 2009. A replication clock for Mycobacterium tuberculosis. Nat Med 15:211–214.

27. Gillet-Markowska A, Louvel G, Fischer G. 2015. bz-rates: A web tool to estimate mutation rates from fluctuation analysis. G3 Genes Genomes Genet 5:2323–2327.

28. Gröschel MI, Owens M, Freschi L, Vargas R, Marin MG, Phelan J, Iqbal Z, Dixit A, Farhat MR. 2021. GenTB: A user-friendly genome-based predictor for tuberculosis resistance powered by machine learning. Genome Med 13:1–14.

29. Guinn KM, Hickey MJ, Mathur SK, Zakel KL, Grotzke JE, Lewinsohn DM, Smith S, Sherman DR. 2004. Individual RD1-region genes are required for export of ESAT- 6/CFP-10 and for virulence of Mycobacterium tuberculosis. Mol Microbiol 51:359– 370.

30. Holt KE, McAdam P, Thai PVK, Thuong NTT, Ha DTM, Lan NN, Lan NH, Nhu NTQ, Hai HT, Ha VTN. 2018. Frequent transmission of the Mycobacterium tuberculosis Beijing lineage and positive selection for the EsxW Beijing variant in Vietnam. Nat Genet 50:849–849.

31. Hsu T, Hingley-Wilson SM, Chen B, Chen M, Dai AZ, Morin PM, Marks CB, Padiyar J, Goulding C, Gingery M. 2003. The primary mechanism of attenuation of bacillus Calmette–Guerin is a loss of secreted lytic function required for invasion of lung interstitial tissue. Proc Natl Acad Sci 100:12420–12425.

32. Hunter JD. 2007. Matplotlib: A 2D graphics environment. Comput Sci Eng 9:90–95.

33. Ioerger TR, O’Malley T, Liao R, Guinn KM, Hickey MJ, Mohaideen N, Murphy KC, Boshoff HI, Mizrahi V, Rubin EJ. 2013. Identification of new drug targets and resistance mechanisms in Mycobacterium tuberculosis. PloS One 8:e75245.

34. Kalyaanamoorthy S, Minh BQ, Wong TK, Von Haeseler A, Jermiin LS. 2017. ModelFinder: fast model selection for accurate phylogenetic estimates. Nat Methods 14:587–589.

35. Kim J-H, O’Brien KM, Sharma R, Boshoff HI, Rehren G, Chakraborty S, Wallach JB, Monteleone M, Wilson DJ, Aldrich CC. 2013. A genetic strategy to identify targets for the development of drugs that prevent bacterial persistence. Proc Natl Acad Sci 110:19095–19100.

36. Kirubakar G, Schäfer H, Rickerts V, Schwarz C, Lewin A. 2020. Mutation on lysX from Mycobacterium avium hominissuis impacts the host–pathogen interaction and virulence phenotype. Virulence 11:132–144.

37. Lewis KN, Liao R, Guinn KM, Hickey MJ, Smith S, Behr MA, Sherman DR. 2003. Deletion of RD1 from Mycobacterium tuberculosis mimics bacille Calmette-Guerin attenuation. J Infect Dis 187:117–123.

38. Li H, Durbin R. 2009. Fast and accurate short read alignment with Burrows--Wheeler transform. Bioinformatics 25:1754–1760.

39. Li H, Handsaker B, Wysoker A, Fennell T, Ruan J, Homer N, Marth G, Abecasis G, Durbin R. 2009. The sequence alignment/map format and SAMtools. Bioinformatics 25:2078–2079.

40. Lim ZL, Drever K, Dhar N, Cole ST, Chen JM. 2022. Mycobacterium tuberculosis EspK Has Active but Distinct Roles in the Secretion of EsxA and EspB. J Bacteriol e00060–22.

41. Love MI, Huber W, Anders S. 2014. Moderated estimation of fold change and dispersion for RNA-seq data with DESeq2. Genome Biol 15:1–21.

42. Manson AL, Cohen KA, Abeel T, Desjardins CA, Armstrong DT, Barry CE, Brand J, Chapman SB, Cho S-N, Gabrielian A. 2017. Genomic analysis of globally diverse Mycobacterium tuberculosis strains provides insights into the emergence and spread of multidrug resistance. Nat Genet 49:395–402.

43. Marin M, Vargas R, Harris M, Jeffrey B, Epperson LE, Durbin D, Strong M, Salfinger M, Iqbal Z, Akhundova I. 2022. Benchmarking the empirical accuracy of short-read sequencing across the M. tuberculosis genome. Bioinformatics.

44. McKinney W. 2010. Data structures for statistical computing in python. Proc 9th Python Sci Conf 445:51–56.

45. Menardo F, Duchêne S, Brites D, Gagneux S. 2019. The molecular clock of Mycobacterium tuberculosis. PLoS Pathog 15:e1008067.

46. Montoya-Rosales A, Provvedi R, Torres-Juarez F, Enciso-Moreno JA, Hernandez-Pando R, Manganelli R, Rivas-Santiago B. 2017. lysX gene is differentially expressed among Mycobacterium tuberculosis strains with different levels of virulence. Tuberculosis 106:106–117.

47. Murphy KC. 2021. Oligo-Mediated Recombineering and its Use for Making SNPs, Knockouts, Insertions, and Fusions in Mycobacterium tuberculosis. Mycobacterial Protocols. Springer. pp. 301–321.

48. Ngabonziza JCS, Loiseau C, Marceau M, Jouet A, Menardo F, Tzfadia O, Antoine R, Niyigena EB, Mulders W, Fissette K. 2020. A sister lineage of the Mycobacterium tuberculosis complex discovered in the African Great Lakes region. Nat Commun 11:1–11.

49. Nguyen L-T, Schmidt HA, Von Haeseler A, Minh BQ. 2015. IQ-TREE: a fast and effective stochastic algorithm for estimating maximum-likelihood phylogenies. Mol Biol Evol 32:268–274.

50. O’Neill MB, Shockey A, Zarley A, Aylward W, Eldholm V, Kitchen A, Pepperell CS. 2019. Lineage specific histories of Mycobacterium tuberculosis dispersal in Africa and Eurasia. Mol Ecol 28:3241–3256.

51. Overbeek R, Olson R, Pusch GD, Olsen GJ, Davis JJ, Disz T, Edwards RA, Gerdes S, Parrello B, Shukla M. 2013. The SEED and the Rapid Annotation of microbial genomes using Subsystems Technology (RAST). Nucleic Acids Res 42:D206–D214.

52. Pandey AK, Yang Y, Jiang Z, Fortune SM, Coulombe F, Behr MA, Fitzgerald KA, Sassetti CM, Kelliher MA. 2009. NOD2, RIP2 and IRF5 play a critical role in the type I interferon response to Mycobacterium tuberculosis. PLoS Pathog 5:e1000500.

53. Pedregosa F, Varoquaux G, Gramfort A, Michel V, Thirion B, Grisel O, Blondel M, Prettenhofer P, Weiss R, Dubourg V. 2011. Scikit-learn: Machine learning in Python. J Mach Learn Res 12:2825–2830.

54. Pepperell C, Hoeppner VH, Lipatov M, Wobeser W, Schoolnik GK, Feldman MW. 2010. Bacterial genetic signatures of human social phenomena among M. tuberculosis from an Aboriginal Canadian population. Mol Biol Evol 27:427–440.

55. Pepperell CS, Casto AM, Kitchen A, Granka JM, Cornejo OE, Holmes EC, Birren B, Galagan J, Feldman MW. 2013. The role of selection in shaping diversity of natural M. tuberculosis populations. PLoS Pathog 9:e1003543.

56. Pérez F, Granger BE. 2007. IPython: a system for interactive scientific computing. Comput Sci Eng 9:21–29.

57. Phelan JE, Coll F, Bergval I, Anthony RM, Warren R, Sampson SL, van Pittius NCG, Glynn JR, Crampin AC, Alves A, Others. 2016. Recombination in pe/ppe genes contributes to genetic variation in Mycobacterium tuberculosis lineages. BMC Genomics 17:151.

58. Safi H, Gopal P, Lingaraju S, Ma S, Levine C, Dartois V, Yee M, Li L, Blanc L, Liang H-PH. 2019. Phase variation in Mycobacterium tuberculosis glpK produces transiently heritable drug tolerance. Proc Natl Acad Sci 116:19665–19674.

59. Sassetti CM, Boyd DH, Rubin EJ. 2003. Genes required for mycobacterial growth defined by high density mutagenesis. Mol Microbiol 48:77–84.

60. Sassetti CM, Rubin EJ. 2003. Genetic requirements for mycobacterial survival during infection. Proc Natl Acad Sci 100:12989–12994.

61. Schmieder R, Edwards R. 2011. Quality control and preprocessing of metagenomic datasets. Bioinformatics 27:863–864.

62. Seabold S, Perktold J. 2010. Statsmodels: Econometric and statistical modeling with python. Proc 9th Python Sci Conf 57:61.

63. Shell SS, Wang J, Lapierre P, Mir M, Chase MR, Pyle MM, Gawande R, Ahmad R, Sarracino DA, Ioerger TR. 2015. Leaderless transcripts and small proteins are common features of the mycobacterial translational landscape. PLoS Genet 11:e1005641.

64. Stanley SA, Johndrow JE, Manzanillo P, Cox JS. 2007. The Type I IFN response to infection with Mycobacterium tuberculosis requires ESX-1-mediated secretion and contributes to pathogenesis. J Immunol 178:3143–3152.

65. Stanley SA, Raghavan S, Hwang WW, Cox JS. 2003. Acute infection and macrophage subversion by Mycobacterium tuberculosis require a specialized secretion system. Proc Natl Acad Sci 100:13001–13006.

66. Tak U, Dokland T, Niederweis M. 2021. Pore-forming Esx proteins mediate toxin secretion by Mycobacterium tuberculosis. Nat Commun 12:1–17.

67. Torres Ortiz A, Coronel J, Vidal JR, Bonilla C, Moore DA, Gilman RH, Balloux F, Kon OM, Didelot X, Grandjean L. 2021. Genomic signatures of pre-resistance in Mycobacterium tuberculosis. Nat Commun 12:1–13.

68. Uplekar S, Heym B, Friocourt V, Rougemont J, Cole ST. 2011. Comparative genomics of esx genes from clinical isolates of Mycobacterium tuberculosis provides evidence for gene conversion and epitope variation. Infect Immun IAI--05344.

69. Van Der Walt S, Colbert SC, Varoquaux G. 2011. The NumPy array: a structure for efficient numerical computation. Comput Sci Eng 13:22–30.

70. Van Der Woude MW, Bäumler AJ. 2004. Phase and antigenic variation in bacteria. Clin Microbiol Rev 17:581–611.

71. Vargas R, Farhat MR. 2020. Antibiotic treatment and selection for glpK mutations in patients with active tuberculosis disease. Proc Natl Acad Sci 117:3910–3912.

72. Vargas R, Freschi L, Marin M, Epperson LE, Smith M, Oussenko I, Durbin D, Strong M, Salfinger M, Farhat MR. 2021. In-host population dynamics of Mycobacterium tuberculosis complex during active disease. Elife 10:e61805.

73. Virtanen P, Gommers R, Oliphant TE, Haberland M, Reddy T, Cournapeau D, Burovski E, Peterson P, Weckesser W, Bright J. 2020. SciPy 1.0: fundamental algorithms for scientific computing in Python. Nat Methods 1–12.

74. Walker BJ, Abeel T, Shea T, Priest M, Abouelliel A, Sakthikumar S, Cuomo CA, Zeng Q, Wortman J, Young SK, others. 2014. Pilon: an integrated tool for comprehensive microbial variant detection and genome assembly improvement. PloS One 9:e112963–e112963.

75. Walker TM, Ip CLC, Harrell RH, Evans JT, Kapatai G, Dedicoat MJ, Eyre DW, Wilson DJ, Hawkey PM, Crook DW, others. 2013. Whole-genome sequencing to delineate Mycobacterium tuberculosis outbreaks: a retrospective observational study. Lancet Infect Dis 13:137–146.

76. Wood DE, Salzberg SL. 2014. Kraken: ultrafast metagenomic sequence classification using exact alignments. Genome Biol 15:1–12.

77. World Health Organization. 2020. Global Tuberculosis Report. https://apps.who.int/iris/rest/bitstreams/1312164/retrieve

78. Zhou X, Stephens M. 2012. Genome-wide efficient mixed-model analysis for association studies. Nat Genet 44:821–824.

